# Optogenetic entrainment of the septo-hippocampal circuit is state conditional and attenuates spatial accuracy

**DOI:** 10.1101/392050

**Authors:** Philippe R. Mouchati, Gregory L. Holmes, Michelle L. Kloc, Jeremy M. Barry

## Abstract

The manipulation of pattern generators in order to impose temporal organization on target nodes while accounting for dynamic changes in behavior and cognitive demand remains a significant challenge for the use of neurostimulation as a therapeutic treatment option. While perturbation through optogentic stimulation can reveal circuit mechanisms that create and locally integrate temporal organization, it is unclear whether superseding endogenous signals with artificial oscillations would benefit or impede hippocampus-dependent cognition or how cognitive demand might affect artificial septo-hippocampal entrainment. Optogenetic MS stimulation in wild-type rats in 3 conditions showed that septal input is more likely to supersede endogenous hippocampal LFP oscillations when animals are at rest or performing a hippocampus-dependent spatial accuracy task. Stimulation during a hippocampus-independent task, however, resulted in compensatory endogenous oscillations. Although stimulation effects on the inter-spike interval of hippocampal pyramidal cells mirrored task-conditional theta entrainment of the LFP, place field properties were unaffected. Analyses of spatial behavior indicate that optogenetic stimulation can attenuate performance and specific measures of goal zone estimation accuracy but otherwise does not affect the rat’s ability to navigate to the target quadrant. The results suggest that the behavioral effect of temporally organizing the septo-hippocampal circuit relative to an artificial theta signal is limited to the accuracy of the rat’s approximation of the goal zone location. These results have significant implications for the therapeutic use of optogenetic stimulation as a means of attenuating cognitive deficits associated with temporal discoordination.

## Introduction

A fundamental feature of the brain is its dynamic organization in time [1]. The manipulation of pattern generators in order to impose temporal organization on target circuits, while taking into account dynamic changes in behavior and cognitive demand, remains a significant challenge to neurostimulation as a treatment option. In rats, the medial septum (MS) has been proposed to act as a putative pacemaker for theta oscillations in the hippocampus [2, 3] by generating rhythmic disinhibition at the synapses of hippocampal pyramidal neurons via long-range axons from the MS to hippocampal GABAergic neurons [4–8]. The resulting temporal coordination by theta phase influences the discharge probability of neurons across the septo-temporal axis [9–11] and the entorhinal-hippocampal loop [12], thereby underpinning mnemonic and spatial processes [13–20]. Temporal discoordination relative to theta oscillations, due to pharmacological or gene manipulation or as a result of status epilepticus in early life, correlates with impairment to memory [21, 22] and performance deficits in a complex spatial task [23].

Artificial theta band stimulation through selective optogenetic stimulation of GABAergic [24] or glutamatergic [8, 25] components of the septal pacemaker in transgenic mice has been shown to supersede endogenous hippocampal theta frequency. Nonselective optogenetic stimulation of the MS in wild-type rats also supersedes endogenous hippocampal oscillations [26, 27], with the caveat that endogenous and artificial oscillations tend to compete during movement speeds greater than 5 cm/s [27]. Optogenetic stimulation therefore has the potential to address several open questions regarding how and when the septal pacemaker entrains or competes with hippocampal oscillations. To this end, it has not yet been shown how behavioral state and cognitive demands placed on the septo-hippocampal circuit might affect entrainment efficacy. Moreover, whether septo-hippocampal entrainment will ultimately aid or impede hippocampal-dependent cognition also remains unknown. This gap in our understanding precludes our ability to employ stimulation techniques for treating cognitive deficits associated with theta rhythmopathy [28]. We therefore carried out two experiments that titrated behavioral state and cognitive demand during nonselective optogenetic septal stimulation. In Experiment 1 we stimulated during limited mobility and cognitive demand while the rats were at rest in a tall, narrow flowerpot. In this condition there was also less endogenous hippocampal theta to compete with artificial theta input. In Experiment 2 we stimulated during 2 versions of a spatial accuracy behavioral task. Although both tasks involved active navigation, the level of hippocampal demand was varied: one version of the task involved a cued goal zone while the other was uncued. Our results indicate that behavioral and circuit state significantly influences septo-hippocampal entrainment efficacy. While stimulation and entrainment perturb several features of temporal coordination in the septo-hippocampal circuit, particularly during the uncued spatial accuracy task, the spatial code is unaffected. Although rats can navigate to the correct quadrant, spatial accuracy relative to the goal zone is attenuated.

## Methods

### Subject details

Seven male Sprague-Dawley rats (obtained from Charles River, Montreal), 3-8 months old were subjects in this study. Animals were housed individually and maintained on a 12 hour light/dark cycle and 85% of baseline body weight and given free access to water. All procedures were approved by the University of Vermont’s Institutional Animal Care and Use Committee and conducted in accordance with guidelines from the National Institutes of Health.

### Injection Surgery

At approximately 3 months old, 4 animals underwent the surgical procedure for the intracranial injection. Rats were anesthetized with a mixture of 5% inhaled isoflurane in oxygen and placed in a stereotaxic frame. All stereotaxic coordinates were relative to bregma [29]. The skull was exposed and a burr hole was placed in the skull. A Hamilton injection syringe was then used to deliver 1 μl bolus injections of the AAV vector (AAV2-hSyn-hChR2(H134R)-EYFP; 5.7 × 10^12^ virus molecules/ml; UNC Vector Core, Chapel Hill, NC) at a rate of 0.1 μl/min into the vertical limb of the diagonal band of Broca in the medial septum (MS) (AP = 0.7 mm; ML= −1.4 mm) and angled 12° medially. The first injection of 0.15 μl was made at a depth of 7.1 mm from brain surface. The syringe was retracted 3 times at 0.3mm steps with injections of 0.2 μl, 0.25 μl and 0.2 μl at each consecutive step. In the final step, the needle was raised 0.2 mm to a final depth of 6.0 mm and an ultimate injection of 0.2 μl was made. The wound was sutured and the rats were returned to their home cages to recover.

### Optical fiber and hippocampal tetrode implants

Detailed descriptions of the optical fiber preparation can be found in a previous publication and are described in brief here [27]. We used a 200 μm multimode optic fiber (Thorlabs, CFLC230-10; Montreal, Canada) as part of our chronic septal implant. The optical fiber was then glued to a 230 μm ferrule (Thorlabs, CFLC230-10; TP01235931). The percentage transmittance of light through the fiber was tested using 100% blue light transmittance (Spectralynx LED source) and measured by a light meter with a photodiode sensor (Thorlabs; Model PM100D). For testing purposes a 50 μm patch cable was used and only fibers that allowed for at least 70% light transmittance at approximately 0.5 mm from the tip of the optical fiber. The MS implant included an optic fiber with an array of 8 recording electrodes glued to the surface that extended 0.25-0.5 mm from the end of the optic fiber. The custom hippocampal implant consisted of 8 separately drivable tetrodes (32 channel electrode array; ‘BLAK drives’; Grasshopper Machine Works, New Hampshire, USA). Each tetrode was composed of 4 twisted 25 μm diameter nichrome wires (California Fine Wire, CA, USA). Each electrode was connected to custom millimax pins (Neuralynx, Montana, USA).

### Optical probe and electrode array chronic implantation surgery

Approximately 2 weeks post-injection, a second surgery was done in order to chronically implant the custom microdrives into the left hippocampus (+ 3.3 mm ML; - 4.5 AP; - 2.0 DV). The optical/recording ensemble was lowered into the MS along the same path previously taken by the Hamilton injection syringe, with the end of the optical fiber lowered to a final depth of ~ 6.5 mm below brain surface. An illustration of the experimental approach using hippocampal septal and hippocampal implants is shown in Fig. 1A.

**Figure 1:**
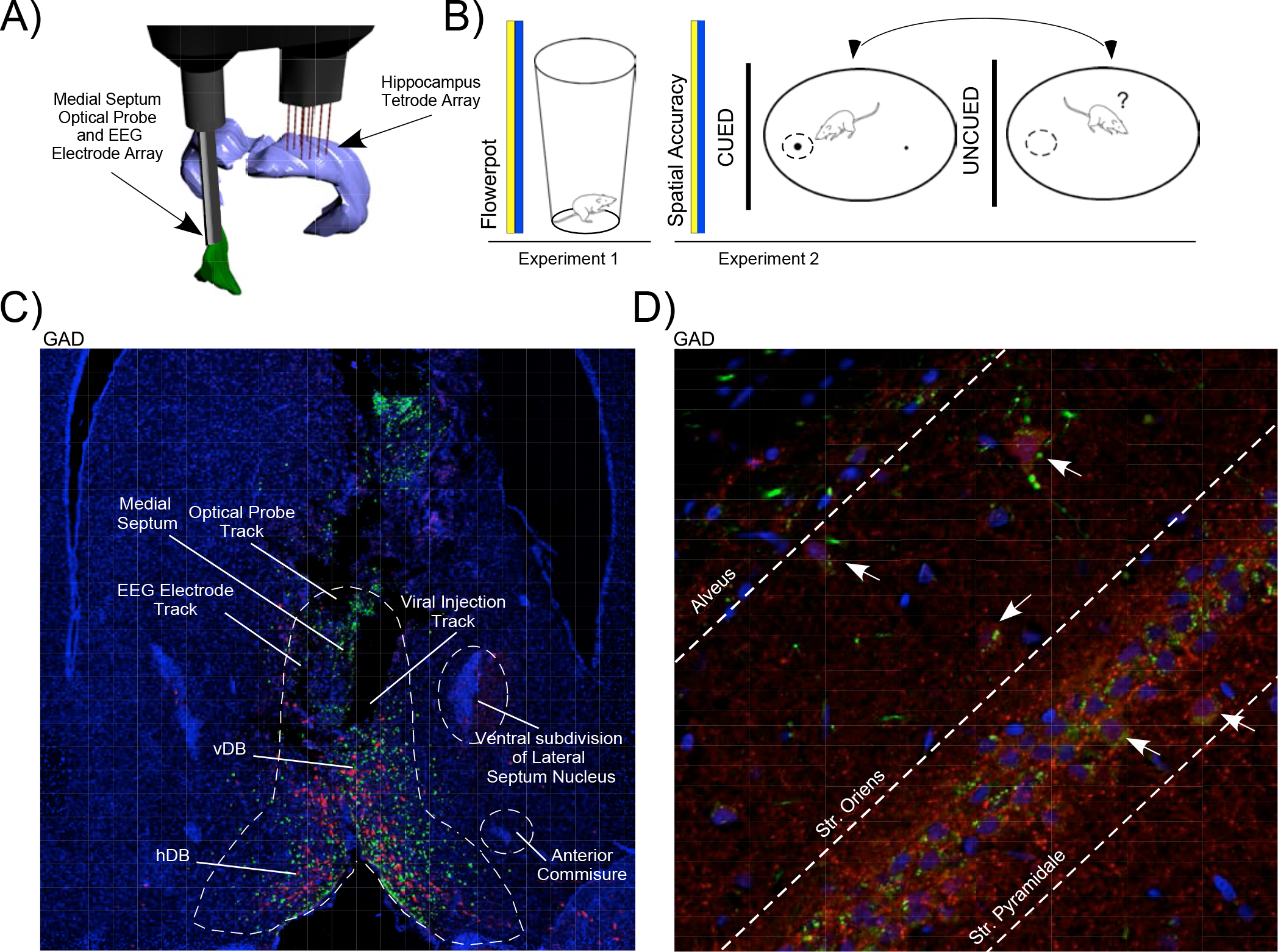
Histology, immunohistochemistry and experimental approach **A)** Experimental approach using optogenetics and electrophysiology in the dorsal hippocampus and MS via custom devices for the delivery of an optical fiber and EEG electrodes in the MS and tetrode array in the hippocampus. **B)** Left - Protocol for Experiment 1 where electrophysiological recordings occurred during 6 min of 6 Hz MS stimulations at both control Y and experimental B stimulation while the rat rests in a tall, narrow flowerpot. Right - Protocol for Experiment 2 where electrophysiological recordings are carried out during 30 min of 6 Hz MS stimulations at both Y and B wavelengths while the rat performs a cued or uncued spatial accuracy task. **C)** Histology of optical and recording probe sites in the MS. Virally transduced MS axons and neurons in green (EYFP), DAPI stained cell nuclei are visible in blue while GABAergic neurons are indicated in red (GAD). Dark areas indicate tracks occupied by the optical probe and EEG wires (Top) and the initial viral injection via the Hamilton syringe (Bottom). These regions are in the center of the MS, flanked by the ventral region of the lateral septum nucleus and the anterior commissure, above the horizontal and vertical limbs of the diagonal band of Broca (hDB and vDB). **D)** Hippocampal GAD expression (white arrows) in red reveals OLM interneurons in str. oriens and basket cells in str. pyramidale as well as virally transduced MS axons (EYFP). DAPI stained cell nuclei are visible in blue. The AAV therefore transduces MS cells whose axonal projections can be traced to the hippocampus, illustrating the circuit and microcircuit levels by which MS stimulation paces hippocampal theta.

Four skull screws (FHC Inc.) were inserted, two were anterior to bregma while the 2 remaining screws were placed over the left and right of the cerebellum. Grounding was achieved via connection to the right cerebellar screw while a reference wire was placed through a small burr hole at brain surface over the cerebellum. Both implants were fixed to the skull via the skull screws and Grip Cement (Dentsply Inc.). The wound was sutured and topical antibiotic applied. The interval between surgery and the beginning of electrophysiological recording was 1 week.

### Stimulation and recording protocols

A 200 μm multimode optic fiber (Thor Laboratories, Budapest, Hungary) was used to connect the implanted optic fiber’s 1.25 ceramic ferrule to Spectralynx, a computer controlled optical LED system (Neuralynx, Montana, USA). The pulse program (Neuralynx, Montana) was used to stimulate the rats with experimental B (wavelength 470 nm) and control Y (wavelength 590 nm). Maximum light intensity was set to 100% (1.8-2.2mW). Light transmittance to the MS via the chronically implanted optical fiber ranged from 70-85%, at the fiber tip. Sine wave stimulatory patterns were generated using the Pulse program (Neuralynx, Montana). The stimulation frequencies of artificial sine waves at 6 Hz (peak/trough = 83.3 msec; period = 166.6 msec) and 10 Hz (peak/trough = 50.6 msec; period = 101.2 msec) were generated by creating an ascending and descending light intensity gradient with a mean light intensity of 52%. The light stimulation intensity was divided into 23 epochs corresponding to 0-255 bits where each increment was a percentage of peak amplitude at 255 bits (i.e., ranging from 1.6 to 100%).

We chose 6 Hz as the primary stimulation frequency as much of the earlier septal stimulation literature noted that hippocampal cells were driven best at frequencies within the 6-8 Hz range [30]. We also wished to shift theta frequency enough from the mean frequency during movement, typically 7.5 Hz, in a manner that would be statistically and functionally distinguishable from baseline endogenous theta oscillations. This frequency would also be more likely to elicit a behavioral effect, as per the experimental hypothesis. We also stimulated at 10 Hz to compare entrainment efficacy between higher and lower limits of the theta band as a function of task condition.

Two experiments were conducted to control for the influence of movement and cognition on the septo-hippocampal circuit and to understand how these variables affect entrainment efficacy through optogenetic MS stimulation (Fig. 1B). Both experiments included optical stimulation in 2 wavelengths, Yellow (590 nm) and Blue (448 nm). The Yellow (Y) wavelength served as control stimulation as this wavelength does not activate channelrhodopsin (ChR2) light-gated ion channels [27]. The Blue (B) wavelength served as experimental stimulation as it causes ChR2 ion channels in the membrane of transduced cells to open. In Experiment 1, stimulation and recording occurred in a flowerpot while the animals were predominantly at rest. Experiment 2 occurred while animals performed 2 versions of a spatial accuracy task. In the 1^st^ version the goal zone was cued and in the 2nd version the goal zone was uncued.

#### Flowerpot

Recording sessions were made while rats rested in a 40 cm high ceramic flowerpot that was 27 cm wide at the base and lined with home cage bedding. The size of the ceramic pot limited movements of the rat to rearing and minor head movements and therefore limited theta oscillations associated with active exploration. The experiment began without stimulation for 2 min, followed by 6 min of 6 Hz Y wavelength stimulation, then 6 min of 6 Hz B wavelength stimulation and ending with an additional 2 min without stimulation.

The flowerpot also served as the location for screening session following the lowering of microdrives. Electrodes were advanced until they contacted the pyramidal cell layer in the CA1 field of the dorsal hippocampus, as characterized by the observation of clustered units as well as 140-200 Hz ripple activity in the local field potentials [23]. Experimental sessions were carried out in the spatial accuracy task when > 20 hippocampal cells were isolated.

#### Cued and uncued spatial accuracy task

Rats began training by exploring a 75 cm diameter circular grey arena with a polarizing white cue card on the wall that covered approximately 90° of arc. The food-restricted rats learned that entry into an invisible 16 cm diameter circular target zone triggered an overhead pellet feeder [31]. The rat’s location in the arena was sampled at 30 Hz (Tracker, Bio-signal Group Corp, Brooklyn, USA) using a firewire digital camera that detected a dark area behind the rat’s head that was made with non-toxic, black permanent marker. When the rat’s position was detected in the goal zone, the computer sent a +5V TTL pulse via the PCI board to a custom feeder in order to trigger the release of a food pellet. For initial training, the center of the target area was “cued” with a white, 20 mm diameter bottle cap. Thirty-min sessions were carried out each day with criterion set to 40 rewards in each of 3 consecutive sessions. After reaching criterion in the cued version of the task, the bottle cap was removed and the rats had to remember the goal location. Criterion was reset to 40 rewards in 3 consecutive sessions. This process of alternating cued and uncued sessions was then repeated while gradually reducing the target zone area from 16 to 8 cm in diameter. Target zone dwell-time thresholds for triggering the feeder also ranged from 0.5 to 1.2 s over the course of training. Taken together, the upper dwell-time threshold and the smallest goal diameter were set so that the rat had to commit to the goal zone. If the rat left the goal zone before the dwell-time threshold was met, the entrance clock was reset and the rat would have to pause in the goal for additional time before reward. Inaccurate performance could then be registered as either increased number of entrances in comparison to the number of rewards or simply as a decrease in the number of rewards. In this manner the rat’s behavior provided a proxy measure of position estimation relative to the goal zone location. A refractory reward period of 5 sec was also set to encourage the animal to leave the target area and forage, thereby spatially sampling the entire arena.

Between cued and uncued sessions rats were placed in the flowerpot alongside the arena for approximately 5 min. During this time the floor panel was cleaned with soap and water and rotated to remove potential olfactory cues. The white cue card remained fixed in relation to the room frame between sessions. Similar to cued and uncued versions of the Morris water maze, cued spatial accuracy is considered to be hippocampus-independent while the uncued version is considered to be hippocampus-dependent [32].

After meeting criterion with goal diameter at 8 cm, the animals underwent the septal injection of the AAV2 virus expressing ChR2 and given a week to recover. The rats were then implanted with the optical probe and electrode array described in Fig. 1A. Screening sessions began after an additional week of recovery. Rats underwent additional training following this final recovery period to ensure that they were still performing at criterion levels and that the surgical procedures or electrophysiological and optical tethers did not interfere with spatial accuracy behavior. The only difference between these training sessions and the initial training period was that detection of the LED on the pre-amplifier was now used to trigger the pellet feeder. Each rat experienced a series of sessions that began with open-loop 6 Hz Y stimulation for 30 min during the cued version of the task. As in pre-implantation training, the rats were not interfered with by the experimenter for this 30 min period. At the end of this session the rats were placed in the flowerpot alongside the arena while the arena platform was cleaned and rotated. The rats were then returned to the arena for another 30 min of 6 Hz Y stimulation during the uncued version of the task. The rats were then returned to their home cages. After approximately 1 hour, the rats were placed back in the arena for successive 30 min cued and uncued sessions with open loop 6 Hz B stimulation in each session. The rats were again placed in the flowerpot between these stimulation sessions for approximately 10 min to allow for cleaning and rotation of the arena platform.

For each rat, the series of 6 Hz stimulation sessions, Yellow Cued (YC), Yellow Uncued (YUC), Blue Cued (BC) and Blue Uncued (BUC) was completed 3-4 times. In each series, the order of cued and uncued sessions during B stimulation was altered at least once in order to test for an effect of trial order. An additional series of 10 Hz stimulation sessions was also completed to compare the resulting electrophysiological effects with the 6 Hz stimulation protocol. Only one stimulation protocol (6 or 10 Hz) was used per day.

The spatial accuracy task allows for several behavioral metrics that reflect the animal’s ability to self-localize relative to the goal zone. Basic measures included the number of triggered rewards, the number of goal zone entrances, and the percentage of session time spent in each arena quadrant (Target, Clockwise, Counter-Clockwise and Opposite). We also utilized the complementary measures of dwell-time and speed relative to the goal zone center to both graphically depict the animal’s choice behavior and calculate search parameters independent of reward. This was calculated using convolution windows (Matlab, Mathworks Inc.) to find continuous bouts of movement ≤ 3 cm/s between a lower time limit of 1.2 s and an upper time limit of 20 s. These speed boundaries were selected so as to distinguish pauses in the goal vicinity from instances where the rat might be travelling through the target quadrant while searching for food pellets. The mean distance of each slow movement epoch from the target center was then calculated and served as a representative measure of goal search accuracy within the target quadrant.

#### Foraging

Three additional animals were used as non-stimulated controls. As in previous experiments [33], rats were food restricted to 80% of baseline body weight and exposed to the same arena used in the spatial accuracy experiments. Food pellets fell at regular 30 sec intervals from the overhead feeder. Each exposure lasted an hour a day for approximately 5 days. By the end of the foraging training, rats sampled the entire surface area of the arena within 30 min. Screening and recording sessions began a week after implantation of the same tetrode array as animals in the other conditions. During screening, electrodes were advanced until they contacted the pyramidal cell layer in the CA1 field, as characterized by the observation of clustered units as well as 140-200 Hz ripple activity in the local field potentials [23]. Recordings consisted of two consecutive 30 min foraging sessions. Between sessions rats were held in the flowerpot for approximately 5 min, in which time the arena floor was cleaned. We wished to determine how successive sessions in the recording environment would affect the properties of theta oscillations. A further question was how movement and the absence of cognitive demand would affect resultant theta phase population vectors. Phase preference results could then be compared between the foraging task and cued version of the spatial accuracy task, given that neither task requires hippocampal-dependent cognition. Results of a previous study showed that foraging does not lead to tight temporal control of CA1 cell activity by local theta oscillations [23].

### Position tracking

The rat’s location in the arena was sampled using a digital camera that detected a light emitting diode (LED) placed near the back of the animal’s head. This tracking information was filtered and recorded utilizing custom software (Neuralynx, Montana, USA) that allows for the synchronization of the rat’s position and speed with properties of the recorded cell and EEG signals. The rat’s behavior and positional information in relation to the goal zone was monitored using the Bio-signal Tracker program (Bio-signal Group Corp, Brooklyn, USA). This program detected the LED in the goal zone and triggered the release of food pellets.

### Signal processing

Rats were tethered to an electrophysiology cable during recording sessions in all conditions. Signals were pre-amplified X 1 at the headstage and channeled through the tether cable to the signal amplifiers and computer interface. Sampling frequency of LFPs (Local Field Potentials) was at 30.3 KHz and filtered at 1-9000 Hz (Neuralynx, Montana), subsampled offline at 3000 Hz. LFP’s were referenced against a 50 μm diameter stainless steel wire (California Fine Wire, CA, USA) placed at brain surface over the cerebellum. LFPs were processed offline using custom software that utilized the Matlab signal processing toolbox ‘spectrogram’ function that returns the time-dependent Fourier transform (FFT) for a sequence using a sliding window (window = 1 sec, overlap = 0.5 sec). This form of the Fourier transform, also known as the short-time Fourier transform (STFT), produces a matrix of frequency, time and signal amplitude in the theta band at 5-12 Hz. The absolute value of the squared FFT signal, or power density, was then converted to decibels. As the amplitude of the signal is therefore a power ratio, it is described in the results using arbitrary units (A.U.).

#### Frequency

Mean frequency in the theta band was calculated for the flowerpot recordings during each 6 min epoch of Y and B stimulation in Experiment 1 as well as in each Y and B condition during performance of the cued and uncued spatial accuracy task in Experiment 2. Peak frequency was determined at the frequency of the maximum amplitude of mean spectral signal during both slow movement speeds (≤ 2 cm/s) and fast movement speeds (between 5 and 50 cm/s).

#### Amplitude

Mean amplitude in the theta band was calculated during each 6 min epoch of Y and B stimulation in Experiment 1 as well as in each Y and B condition during performance of the cued and uncued spatial accuracy task in Experiment 2. Amplitude normalization was carried out by dividing the spectral signal over its sum. Peak amplitude was determined by the maximum value of the non-normalized speed-sorted mean spectral signal.

#### Speed/Theta properties

As in previous work [27, 34], we analyzed the linear relationship between animal speed and theta band frequency and amplitude during each version of the spatial accuracy task. We also analyzed speed and theta frequency during foraging sessions in non-stimulated control animals.

#### Unit isolation and classification

Single units were sampled at 30.3 KHz and filtered at 300-6000 Hz (Neuralynx, Montana). All tetrode signals were referenced against a tetrode wire near the corpus callosum. The activity of individual units was separated offline into different clusters based on their waveform properties. Waveform properties were defined in 3-dimensional feature space using custom spike-sorting software (Plexon, Dallas, TX). The primary quantitative measure of cluster quality provided by OFS (Offline Sorter) was J3, a non-parametric measure of the quality of cluster sorting [35]. A measure of the average distance between unit clusters (J2) is first calculated: J2 = ΣN(u)E(m(u)-m). The summation is over units u, N(u) is the number of points in unit u, E(x) represents the Euclidean distance squared. m(u) is the cluster center for unit u, and m is the grand center of all points in all units. The final measurement, J3 is then calculated by dividing J2 by a 3^rd^ variable (J1): J1 = ΣΣE(f(u,i)-m(u)). This is a measure of the average distance in 3-dimensional feature space between points in a cluster (f) from their center (m). E(x) represents the Euclidean distance squared (**x**̃ * x). The summations are over units u, and over feature vectors (points) in each unit i. J3 therefore takes on a maximum value for compact, well-separated clusters.

Finally, spikes that had a refractory period of less than 1 msec and units that were not determined to be pyramidal cells or interneurons [36, 37] were removed from analysis. Putative OLM and basket cell activity were distinguished via estimated recording depth, theta amplitude, phase preference[38] and phase offset between recording sites to distinguish between str. oriens and str. pyramidale. Pyramidal cells, interneurons and axonal activity were distinguished from each other via activity and waveform properties as described previously [36, 37, 39] but are briefly described here. Interneurons discharge at high rates (> 10 Hz), have short duration waveforms (peak to trough duration ~ 250 μsec), never show complex spike bursts and tend not to exhibit circumscribed firing fields. Pyramidal cells discharge at lower rates (< 9 Hz), have wider waveforms (peak to trough duration ~500 μsec) and fire in complex-spike bursts. Pyramidal cells may act as “place cells” whose activity is typically confined to a small, cell-specific region called the “firing field” [33, 40]. Axonal activity typically exhibits a short duration waveform (peak to trough duration < 250 μsec) with a triphasic waveform shape [36, 39].

#### Autocorrelation and corresponding periodicity

Spike times were binned at 10 msec intervals for each cell during flowerpot recordings and during spatial accuracy recordings in each stimulation and task condition. Autocorrelations between spike times [41] were computed via the Matlab cross-correlation function (‘xcorr’) that measures the similarity between the vector and the shifted (lagged) copies of the vector as a function of the lag. Autocorrelation histograms were plotted based on these autocorrelations that not only illustrated changes in the temporal organization of inter-spike intervals but any potential change in the number of events that might be caused by B stimulation. A variation of the cross-correlation function (‘coeff’) was used to generate correlations equal to 1 at lag times equal to 0. The resulting autocorrelogram (ACG) was then used to calculate the modulation periodicity between the peak correlation times after time 0. Modulation frequency was determined through the reciprocal of the periodicity of these peak correlation times. The magnitude of the largest peak and the largest trough after time 0 were then subtracted to provide a measure of theta modulation depth (Supp. Fig.1). As modulation frequencies tended to be overestimated or inaccurate when modulation depth was low, frequencies were analyzed with modulation depth as a covariate using GEE (General Estimating Equations). Although peak autocorrelation values were not used in analysis, they are provided in several examples as a reference for modulation depth. Cell data recorded during spatial accuracy experiments were subject to speed filtering. Spikes occurring during epochs of animal movements ≥ 2 cm/s were excluded from analysis.

#### Place cell firing fields

Several firing field properties were analyzed to determine the effect of task and stimulation condition on the spatial tuning of hippocampal cells as well as to verify whether or not each unit met place cell criteria [33]:

*Firing rate* - The number of action potentials fired by the cell in the firing field divided by dwell-time in the field. The mean overall firing rate was also determined by dividing the total number of action potentials by the length of the recording session.
Firing field coherence - Coherence was calculated as previously described [33] and estimates the regularity of the firing rate transition from pixel to pixel in relation to the spatial firing distribution. Cells with field coherence values > 0.2 were considered to be place cells.
*Field size* - The size of firing fields considered for analysis was set to a lower limit of 9 contiguous pixels and a maximum of 700 pixels (approximately 60% of the arena surface).
*Firing field stability* - Positional firing stability was estimated from the similarity of the place cell’s spatial firing pattern between each stimulation and task condition during spatial accuracy performance. In order to test for the influence of task and stimulation, direct comparisons were made between Y and B stimulation sessions during the uncued task, Y and B stimulation sessions during the cued task and Y stimulation sessions during the cued and uncued task. Similarity is defined as the Fisher z-transform of the product-moment correlation between the firing rates in corresponding pixels for both intervals.

#### Phase preference

The number of action potentials from each cell relative to the phase of the theta band in the local field potential was analyzed via phase preference analysis as in previous studies [23]. Several criteria were imposed on the selection reference channel that served as the representative LFP signal for each tetrode. LFPs were only considered if they exhibited 140-200 Hz ripple activity, when the rat was at rest, and the wire had at least one cell. The selected LFP was filtered between 4-14 Hz using Chebyshev type 2 filters, then the phase was extracted using the Hilbert transform. All the spikes recorded from this tetrode and from neighboring tetrodes were assigned a phase value from the ongoing theta phase from the referenced LFP. The trough of the theta cycle was assigned to 0°/360° while the peak of the theta cycle was assigned to 180°. The mean preferred phase of firing was calculated for each cell by averaging the circular phase angle of each spike divided by the total spike count. Cells were considered to exhibit a significant phase preference when the p-value calculated with Rayleigh’s test for non-uniformity of circular data [46] was determined at p ≤ 0.01. Rayleigh’s test for non-uniformity of circular data was calculated as [*Rbar* = *n*\**r*] [42] where n is the sum of the number of incidences in cases of binned angle data and r is the resultant vector length of the distribution. The angle (Dir) and length of the resultant vector (RBAR) for the individual cell and the population of cells for each rat were then calculated and plotted for each stimulation condition in both experiments. As a compliment to RBAR, we also measured population dispersion levels [δ =1 − T_2_ / (2 * RBAR^2^)] [42] and measured the variability of the population vector relative to theta cycle. If the population RBAR levels are low, dispersion levels should be high.

To test the changes in the circular distribution of preferred firing phase between stimulation conditions in experiment 1, we used the parametric Watson-Williams multi-sample test for equal means, used as a one-way analysis of variance for circular data [43].

### Statistical Analyses

As the behavioral, single unit and EEG datasets contain data from multiple sessions in a repeated measures design in single animals, the assumptions of independence of observations are invalid. The observations for each of these measures within single animals are likely to be correlated and these data can be represented as a cluster. In this case, the existence of a relationship between each measure of interest within an individual animal may then be assumed [44]. We therefore used GEE (General Estimating Equations; SPSS; Armonk, NY), a class of regression marginal model, for exploring multivariable relationships between clustered signal property data or behavioral measures for individual animals sorted by stimulation condition. Unless otherwise indicated, analyses are by rat. The model was adjusted according to the distribution of each analyzed variable, i.e., gamma with log link models were used for non-normally distributed data while poisson loglinear distributions were used for count data. Importantly, in both Experiment 1 and Experiment 2 we used a repeated measures design using the same animals between stimulation conditions to compare the effects of Y or B light stimulation or task conditions on electrophysiological and behavioral measures. In Experiment 2, the principal condition used as a reference for comparison across stimulation conditions was the BUC session. When warranted, the YC baseline stimulation condition was also used as a reference.

### Histological procedures

Rats received a lethal dose of isoflurane and were perfused intracardially with saline followed by 4% formaldehyde. Brains were extracted and stored in 4% formaldehyde. Frozen coronal sections (30 μm) were cut and stained with cresyl violet. Electrode positions in the hippocampus (Supp. Fig. 2A), optical probe and electrode positions in the MS (Supp. Fig. 2B) were assessed in all rats. The volume of the EGFP-labeled viral expression area in the MS was calculated using previously described methods [24]. Coronal brain slices (40 μm thick) containing the MS area for each rat was stained with DAPI and tiled images were taken on a Nikon C2 Confocal Microscopy System (Nikon Corporation, Tokyo, Japan) using a 10X (0.45 NA) objective. All images were obtained using identical parameters and dimensions for consistency in the analysis. Images of the MS were obtained along the AP axis close to the beginning, center, and terminus of the EGFP expression area to calculate the volume of expression. Using ImageJ (FIJI), tile images were stitched together [45] and a region of interest (ROI) was drawn around the medial septum and the diagonal band of Broca (ROI area = 1.47 mm^2^); the same ROI was used across animals. For each stitched image (3 per animal), EGFP pixel area was analyzed within the ROI. The final viral expression volume was obtained by summing the EGFP area of each stitched imaged and multiplying by the distance between imaged slices.

Septal sections were stained with anti-choline acetyltransferase (polyclonal rabbit anti-ChAT antibody, Synaptic Systems), anti-glutamate decarboxylase 2 (polyclonal guinea pig anti-GAD 2, Synaptic Systems), and anti-Vglut2 (polyclonal chicken, Synaptic Systems). All imaging was performed in SPOT 5.0 advanced imaging software (SPOT Imaging Solutions, Michigan, USA) using a Nikon DS-Fi1 color digital camera on a Nikon E400 microscope with FITC, TRITC, and DAPI filter set. Two 20x images were taken from septal sections of each animal stained for GABAergic, cholinergic and glutamatergic neurons. An additional 40x image was taken of a hippocampal section stained for glutamatergic axon terminals projecting from the MS that expressed VGLUT. Cells were considered co-localized when GAD and ChAT overlapped with GFP. Only cells with distinct nuclei as determined by the presence of DAPI were considered to be representative examples. For VGLUT expression, MS axon projection terminals that expressed GFP and VGLUT in hippocampus were considered to be co-localized.

## Results

We examined the role of the behavioral and septo-hippocampal circuit state on optogenetic entrainment using two experiments. In experiment 1 (N = 3), 6 Hz Y and B MS stimulations were made while animals were in the flowerpot. In this condition, we hypothesized that there would be less competition between endogenous hippocampal theta and artificial theta oscillations without movement or cognitive demand. In experiment 2 (N = 3), animals performed a spatial accuracy task where the goal zone was either cued (C) or uncued (UC). In the cued condition the task is hippocampus-independent while in the uncued condition the task is hippocampus-dependent [32]. Each task was repeated to allow for comparison between continuous open-loop 6 Hz Y and B MS stimulation during each session (Yellow Cued = YC, Yellow Uncued = YUC, Blue Cued = BC, Blue Uncued = BUC). We hypothesized that changes in cognitive demand between the cued and uncued tasks would affect the septo-hippocampal circuit in a manner that would influence the efficacy of hippocampal entrainment. To compare hippocampal temporal phenomena during movement, but in the absence of cognitive demand, we also obtained electrophysiological recordings from an additional 3 animals while foraging for food pellets.

### Confirmation of MS probe and electrode placement and viral transduction volume

Examples of immunohistochemistry of ChR2 transduction in the MS, including co-expression of ChR2 in cholinergic and GABAergic cells, can be seen in Supp. Fig. 2C. As VLGLUT expresses preferentially in axon terminals, we found sparse expression in the soma of MS cells [46]. We therefore show co-localization of VGLUT and GFP at the axon terminals of septal projections in str. pyramidale and str. oriens in CA1 (Right Supp. Fig. 2C). An example of GFP and GAD expression in conjunction with septal probe and electrode array placement in the MS can be found in Fig. 1C. The axonal projections from transduced septal cells to GAD positive interneurons in str. oriens and str. pyramidale layers of CA1 in hippocampus are shown in Fig. 1D. To interpret the effect of optogenetic stimulation on electrophysiological and behavioral results we estimated the volume of viral transduction in the MS (Supp. Fig. 2B). Estimated GFP expression volumes were consistent with the estimated total volume of the adult rodent MS (mean = 1.69 ± 0.3 mm^3^; Supp. Table 1). Notably, despite having the densest GFP expression in the dorsal region of MS, rat P2 lacked viral expression in the more ventral horizontal limb of the diagonal band (DB). Rats P3 and P4 both showed comparatively similar GFP expression density, which was localized in both the medial septum and vertical limb and horizontal limb of the DB.

## Experiment 1

### MS optical stimulation entrains hippocampal theta band oscillations

As illustrated in Fig. 2A, 6 Hz B stimulation in the MS entrains oscillations in the MS and hippocampus. Example spectrograms for Y and B stimulation in the MS and hippocampus for the full 6 min sessions are shown in supplemental material (Supp. Fig. 3). Fig. 2B shows an example of the mean amplitude within the theta band for the MS and hippocampus for the 6 mins of Y and B stimulation while a microcircuit level experimental model by which optical stimulation of the MS influences hippocampal oscillations is provided in Fig. 2C-D. We note that in this condition when the animal is at rest and has limited movement there is little endogenous MS theta.

**Figure 2:**
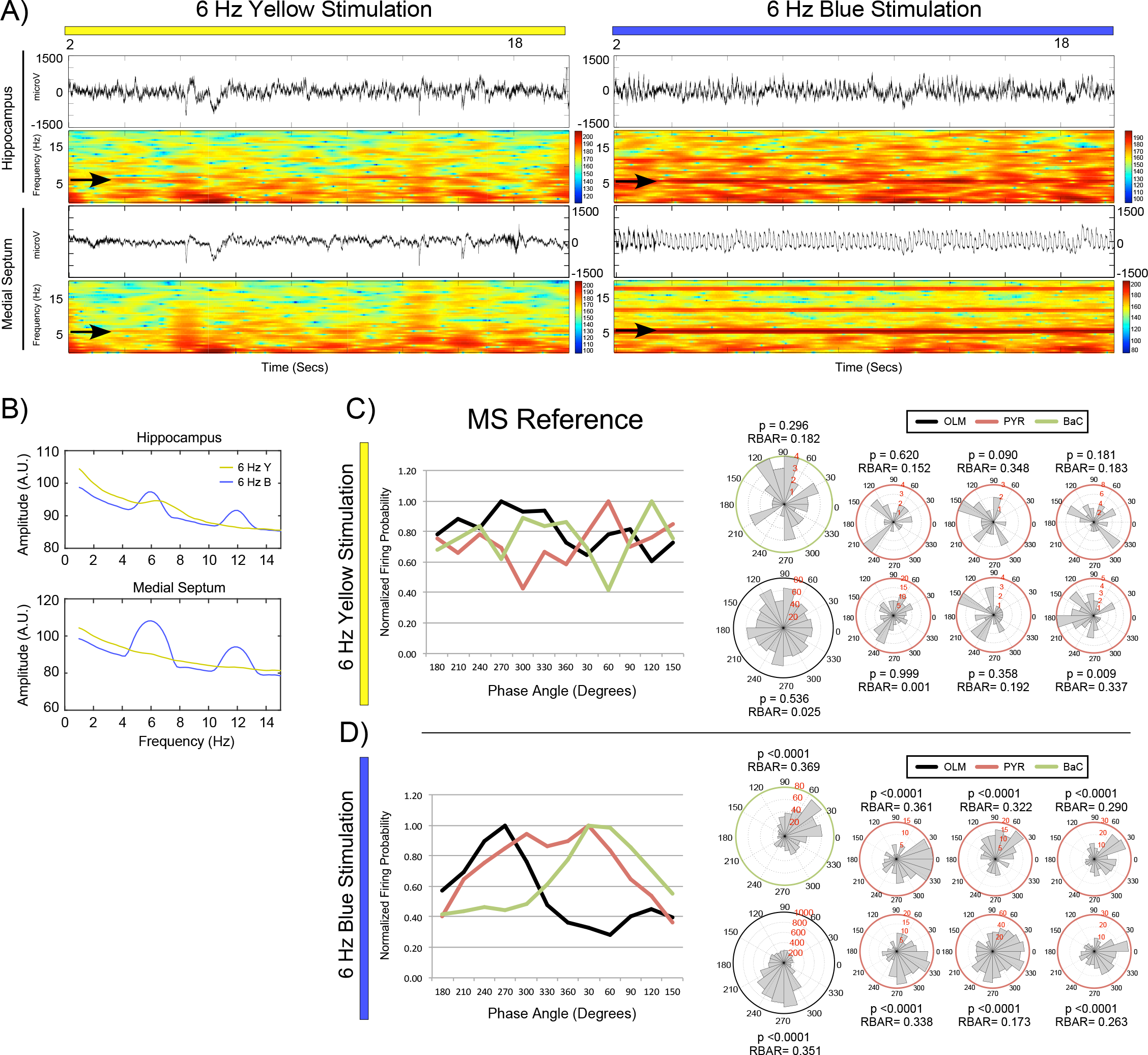
Experiment 1 – Representative results from six min of 6 Hz Y and B MS stimulation and recording in a narrow flowerpot from P3. **A)** Representative results from approximately 20 sec epochs of raw EEG and corresponding spectrogram of CA1 (Top) and MS (Bottom) signals during Y (Left) and B stimulation (Right). Theta band signals are indicated by black arrows. **B)** Amplitude for 6 min Y and B stimulation epochs in hippocampus (Top) and MS (Bottom) suggests peak amplitude shifts to 6 Hz, matching the MS input frequency. **C)** Mean normalized firing probability of simultaneously recorded cells in str. pyramidale, 17 pyramidal cells (PYR in red) and 6 putative basket cells (BaC in green), and 3 putative OLM cells from str. oriens (black) during Y (Top) and B (Bottom) MS stimulation relative to MS theta. During Y stimulation there is little MS theta signal and little temporal structure in the organization hippocampal cell activity as neither cell type exhbits clear peak and valleys of firing probability by theta phase. Firing phase of hippocampal cells is significantly reorganized relative to the septal theta reference during B stimulation. Peak pyramidal cell activity occurs between the peaks of both interneuron types. Top Right - Example rose plot histograms for each MS referenced neuron type during Y (Top) and B (Bottom) MS stimulation. The resultant theta phase vector (RBAR) and significance (p-value) for each cell is shown. Bottom Right - As indicated by firing probability plots in C, interneurons become phase locked to the artificial MS signal. BaC and OLM interneurons are phase locked to opposite phases of MS theta while PYR cells fire between these phases. In this manner, MS stimulation paces the frequency of hippocampal LFP oscillations in the theta bandwidth.

Consequently, there is little temporal organization of hippocampal neurons relative to theta. GEE analysis of B stimulation induced changes to the theta band properties of representative LFP signals from the hippocampus and MS of three animals are shown in Fig. 3A. Mean peak theta frequency decreased significantly between Y (6.4 ± 0.0 Hz) and B (6.0 ± 0.09 Hz) stimulation epochs (p < 0.0001), indicating a shift in peak hippocampal theta oscillations to the septal input frequency. Meanwhile, no significant changes were found relative to Y or B stimulation for hippocampal mean peak theta amplitude or mean normalized theta amplitude. In contrast, significant changes to both theta amplitude and frequency in the MS were caused by B stimulation (Fig. 3B). Mean peak theta frequency increased significantly between Y (5.3 ± 0.22 Hz) and B (6.0 ± 0.0 Hz) stimulation epochs (p = 0.001), indicating a universal shift in peak hippocampal theta frequency in correspondence with the artificial theta band stimulation. B stimulation also caused significant increases in mean peak theta amplitude of the septal signal (p < 0.0001), as well as significant increases in mean normalized theta amplitude (p < 0.0001). It is important to note that both the signal and amplitude of MS theta during baseline strongly suggest that there was a general absence of MS theta oscillations, particularly since the peak theta frequency across animals approached the lower cut-off limit for theta frequency at approximately 5 Hz. Thus, optogenetic MS stimulation can generate artificial theta oscillations in the absence of robust endogenous oscillations while the rats are awake but with limited movement.

**Figure 3:**
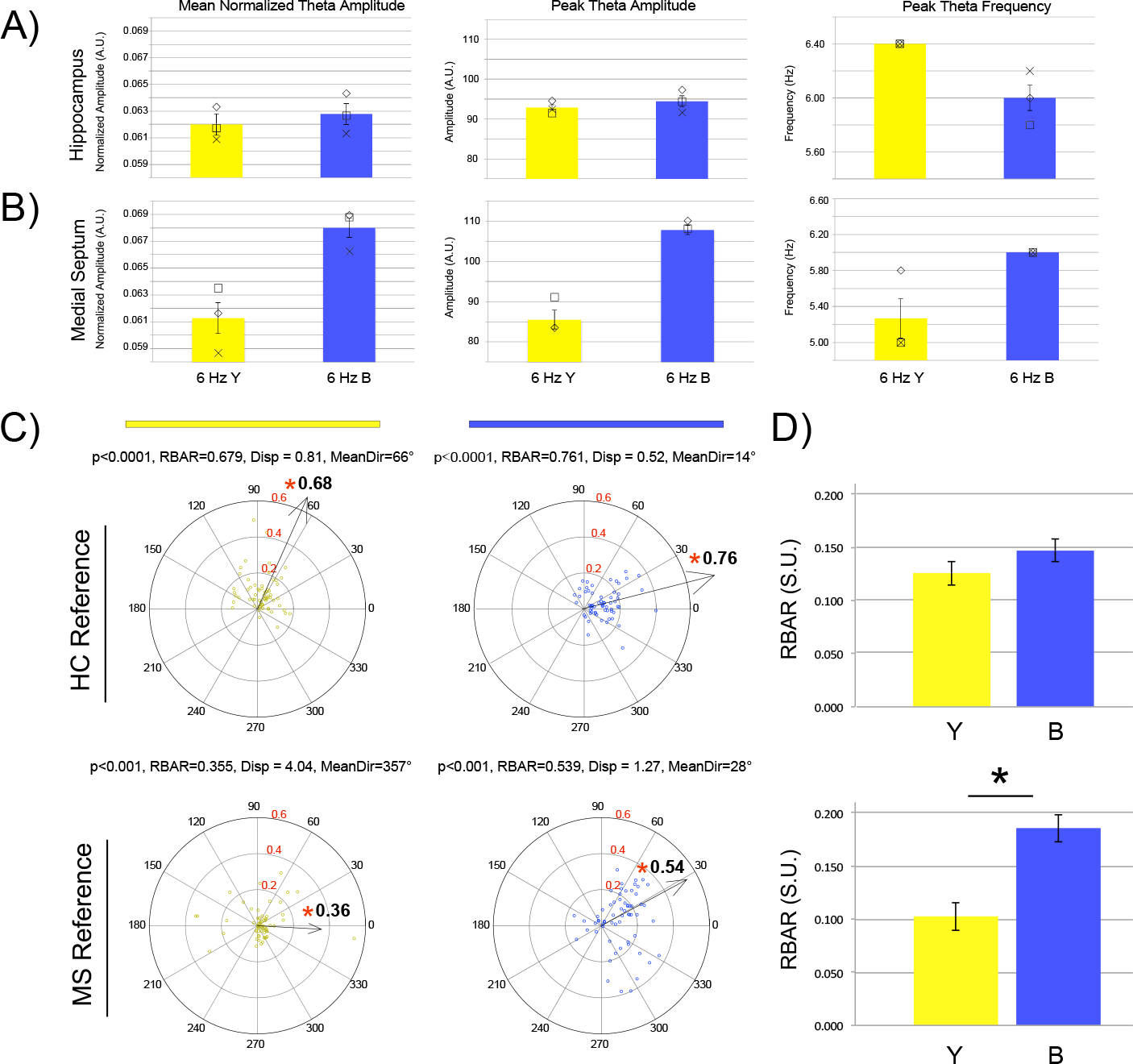
Experiment 1 - Mean and SE of normalized theta amplitude, peak theta amplitude and peak theta frequency in the hippocampus **(A)** and MS **(B)** during 6 Hz Y and B stimulation. Mean scores from individual rats are also shown (P0 = X, P3 = Square, P4 = Diamond). **C)** Phase preference circular statistics for hippocampal CA1 pyramidal cells from all 3 animals (n = 64) during 6 Hz Y (Left) and B (Right) MS stimulation epochs relative to local hippocampal theta (Top) and MS theta (Bottom). Each circle represents the mean angle and magnitude of phase preference for an individual pyramidal cell. The arrow indicates the angle and significance of the resultant population phase vector. RBAR levels for the circular plot are shown in red and mean vector RBAR is presented at the tip of each arrow. A red asterisk denotes significance. During Y stimulation (Left), the hippocampal cell population has a robust mean resultant vector (Arrow) at 66° relative to local theta and a weaker resultant vector at 0° relative to septal theta. During B MS stimulation, the cell phase vectors relative to both references shift into alignment toward the trough of theta (Right). The significance and magnitude of the phase preference (p and RBAR values) as well as the mean direction (MeanDir) and levels of dispersion (Disp) are shown. **D)** Septal stimulation does not change the mean RBAR of each cell relative to local theta but does cause a significant increase relative to the septal signal.

### MS optical stimulation aligns septal and local theta phase preference of hippocampal neurons

The effect of stimulation condition on theta phase preference of CA1 pyramidal cells (n = 64), pooled for all 3 animals (n = 33, 14 and 17), is shown in Fig. 3C. Significantly different mean resultant vector lengths (RBAR) and phase angles were found for the pyramidal cell population in Y and B stimulation conditions referenced to local theta band signals from the hippocampus (Y = 0.679 at 66°; B = 0.761 at 14°) and MS (Y = 0.355 at 357°; B = 0.539 at 28°). Watson-Williams test comparing Y and B stimulation epochs for reveal that theta phase preference was significantly altered for hippocampal cells relative to hippocampal (p < 0.0001) and MS (p = 0.027) theta references. Paired t-test comparisons for the RBAR values for all cells by the LFP in each reference region and stimulation condition demonstrate that B stimulation had no effect on the average length of the resultant phase vector relative to local hippocampal theta (p = 0.115) but did have a significant effect relative to septal theta (p < 0.0001). While the phase organization by the septal reference signal was weak during baseline Y stimulation, it was significantly stronger during B stimulation. The phase angle shift of CA1 pyramidal neurons toward the trough of hippocampal theta, in alignment with the septal signal during B stimulation, suggests a lack of competition between septal inputs and other hippocampal inputs in this condition. Notably, this lack of competition does not result in a perturbation of the magnitude of hippocampal theta phase preference but suggests that the MS dictates the preferred phase angle.

### MS optical stimulation entrains burst-timing of a subset of pyramidal cells

Using ACGs we determined the inter-spike intervals of CA1 pyramidal cells to examine the influence of artificial theta stimulation protocols on the temporal organization of complex bursts. Despite robust entrainment of the LFP and phase preference, only 25% of recorded cells were found to have altered temporal firing patterns between Y and B stimulation, as determined by modulation depth. We refer to this upper quartile of stimulation responsive cells as U25MD. Example autocorrelation histograms of this stimulation responsive subset are shown in Fig. 4A and a histogram describing the modulation depth of the cell population is shown in Fig. 4B. Tukey’s hinge analysis suggested that the cut-off modulation depth value for this responsive subset, above the 75^th^ percentile was > 0.0085 (A.U.). GEE analysis revealed that in comparison to Y stimulation, this subset of cells (U25MD Fig. 4C-E) exhibited a significant increase in modulation depth (p < 0.0001). While this result is a defining feature of this subset, these cells also had a decrease in modulation frequency toward 6 Hz (p < 0.0001). Modulation depth of the remaining 75% of the cells (R75MD in Fig. 4 C-E) was significantly attenuated in response to B stimulation (p < 0.0001) while the modulation frequency increased significantly. However, as ACG analysis tended to over-estimate periodicity when modulation depth was low, this increase in frequency was likely an artifact of decreased modulation. Finally, the U25MD cells have a significantly higher overall mean firing rate during baseline than the R75MD cells (p = 0.032). Importantly, B stimulation did alter the overall mean firing rate in either cell population (U25MD or R75MD) relative to baseline (p = 0.224).

**Figure 4:**
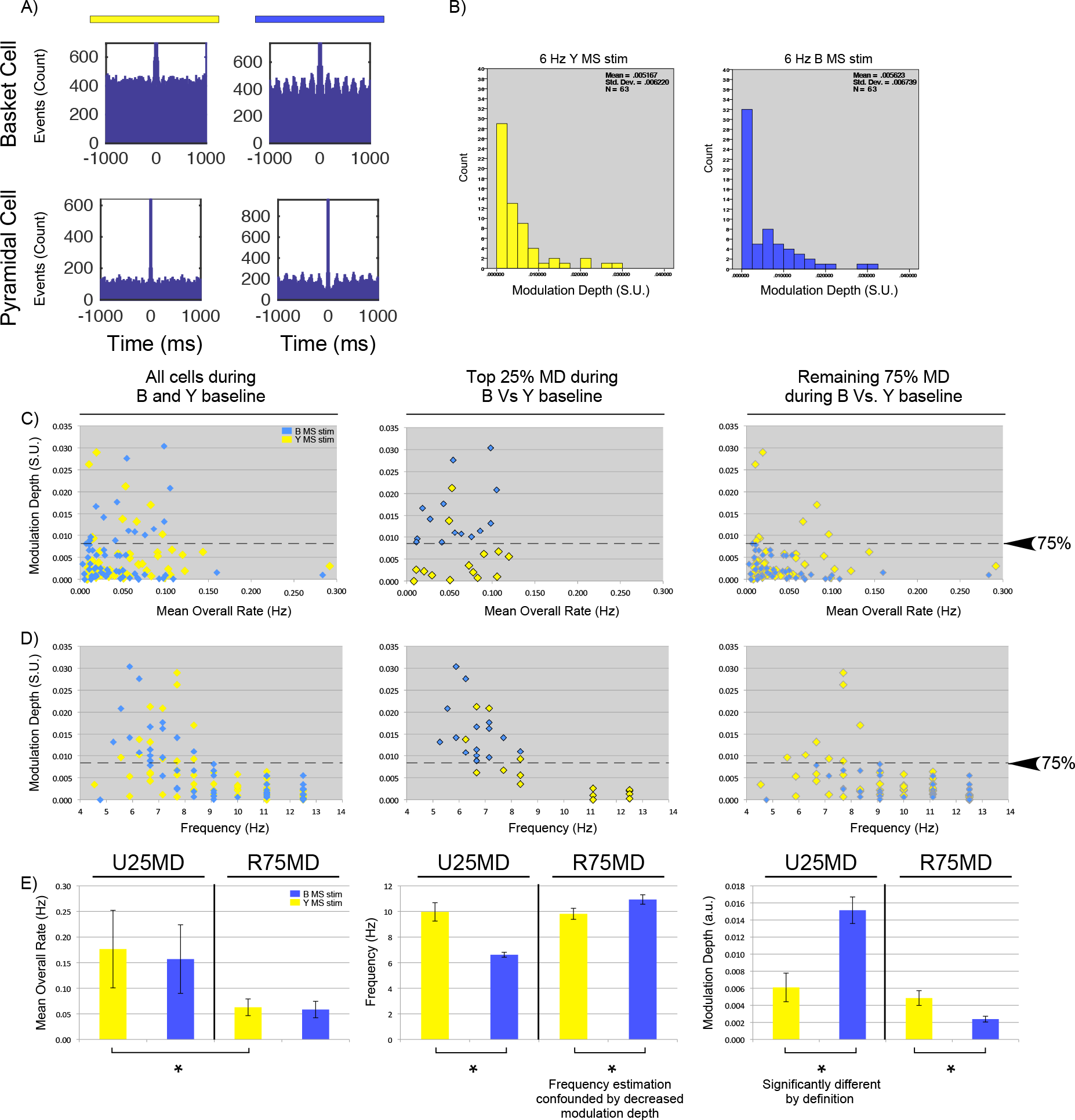
Experiment 1 - Spike timing population dynamics during 6 Hz Y and B MS stimulation. **A)** Example autocorrelation histograms during 6 Hz Y (Left) and B (Right) stimulation for a putative basket cell interneuron (Top) and pyramidal cell (Bottom). B stimulation causes the spike timing of these neurons to shift toward intervals that match the 6 Hz MS input frequency. **B)** Histograms illustrating the lack of overall change in the mean and standard deviation of modulation depth of sampled cells in response to B stimulation, but a change in a subset of cells with a resultant modulation depth > 0.01. **C-D)** Scatter plots of all cells comparing modulation depth against mean overall rate (Left C) and modulation frequency (Left D). The modulation depth during B stimulation was then ranked by quartile and the population split between the top 25% of the most modulated cells (U25MD) and the remaining 75% of the cell population (R75MD) (Middle and Right C-D respectively). Dashed line represents Tukey’s hinge between the top quartile and the remaining 75th percentile. **E)** Mean and standard error for the overall rate, modulation frequency and modulation depth of each cell population (U25MD vs. R75MD) during each stimulation condition. Mean overall rate does not change during B stimulation for either population. U25MD cells tend to have a higher mean baseline-firing rate (Left). These cells are also more likely to shift frequency to match the MS input frequency at 6 Hz (Middle). While the R75MD cells appear to significantly increase frequency, this result is confounded by the decrease in modulation depth for this population that causes an overestimation of frequency. As the defining feature of this subpopulation, modulation depth increases significantly during B MS stimulation (Right).

The main result of this experiment is that optogenetic MS stimulation entrains the frequency of hippocampal synaptic oscillations within the theta bandwidth, shifts the preferred theta phase of pyramidal cells, and influences the burst-timing patterns of approximately a quarter of sampled pyramidal cells. The results also indicate that cells that were most active prior to stimulation were more likely to be entrained by MS stimulation. Cells that were less active prior to stimulation were more likely to exhibit attenuated temporal modulation. Thus, artificial theta stimulation interacts with recent cell activity to recruit active cell assemblies. In summary, septal entrainment of hippocampal oscillations in the absence of robust endogenous theta signals is effective and relatively uncomplicated. In Experiment 2 we address the hypothesis that artificial theta stimulation and endogenous theta oscillations may compete during active navigation and that this competition may be further influenced by levels of cognitive demand.

## Experiment 2

### MS entrainment of hippocampal LFP theta band oscillations is influenced by task and speed

To examine the influence of movement and changes in cognitive demand on both the temporal dynamics of the septo-hippocampal circuit as well as the efficacy of its entrainment by artificial theta stimulation, we conducted a 2^nd^ experiment. This experiment involved movement and active navigation toward a cued goal in one condition and an uncued goal in a second condition, during either control 6 Hz Y or 6 Hz B MS stimulations. In this manner we test for septo-hippocampal entrainment efficacy during movement while simultaneously testing for the influence of cognitive demand by behaviorally titrating spatial cognition in each stimulation condition (YC, YUC, BC and BUC). Previous studies have found that the linear relationship between theta properties and animal speed to be an important correlate with spatial memory performance [34, 47]. As MS stimulation in this study is open loop, entrainment could eliminate any potential biological relevance provided by an endogenous theta signal that changes dynamically with speed. Therefore, we also carried out separate measurements for the linear relationship between animal speed and theta frequency or theta amplitude.

Animal speed has an important effect on theta band properties [34] and prior studies have described optogenetic lateral septum stimulation effects on movement speed [48]. GEE analysis revealed no main effect of stimulation and task condition on mean animal speed (YC = 14.03 ± 1.63 cm/s; YUC = 11.63 ± 1.35 cm/s; BC = 12.63 ± 1.46 cm/s; BUC = 10.69 ± 1.24 cm/s; p = 0.393).

An illustration of the relationship between movement speed and hippocampal theta band properties of frequency and amplitude during each condition are shown in Fig. 5A. Speed-sorted spectrograms for each of these example sessions can be found in supplementary material (Supp. Fig. 4). Y stimulation in each condition did not perturb the expected linear relationship between theta amplitude and frequency in relation to speed. As we previously reported [27], we found that B stimulation during the cued condition at movement speeds ≥ 5 cm/s resulted in a separation of the theta band into two predominant frequencies, 6 Hz and approximately 9 Hz. During slower movement speeds at ≤ 2 cm/s, the hippocampal theta frequency at 6 Hz was more prevalent. Plotting frequency of theta cycles across all speeds during BC suggests that the 9 Hz component of the signal still increases with movement speed but is shifted approximately 1.5 Hz faster than baseline conditions during YC. A further example of this competition between a putative endogenous theta signal and the septal stimulation frequency in relation to movement speed from an additional animal is shown in Supp Fig. 5. Importantly, during the hippocampal-dependent BUC session, theta oscillations at 6 Hz dominated the bandwidth, regardless of movement speed (Fig. 5A). Endogenous theta oscillations were therefore less likely to compete with the septal stimulation frequency in this condition. Accordingly, 6 Hz B stimulation attenuated the relationship between animal speed and theta properties of frequency and amplitude, particularly during BUC.

**Figure 5:**
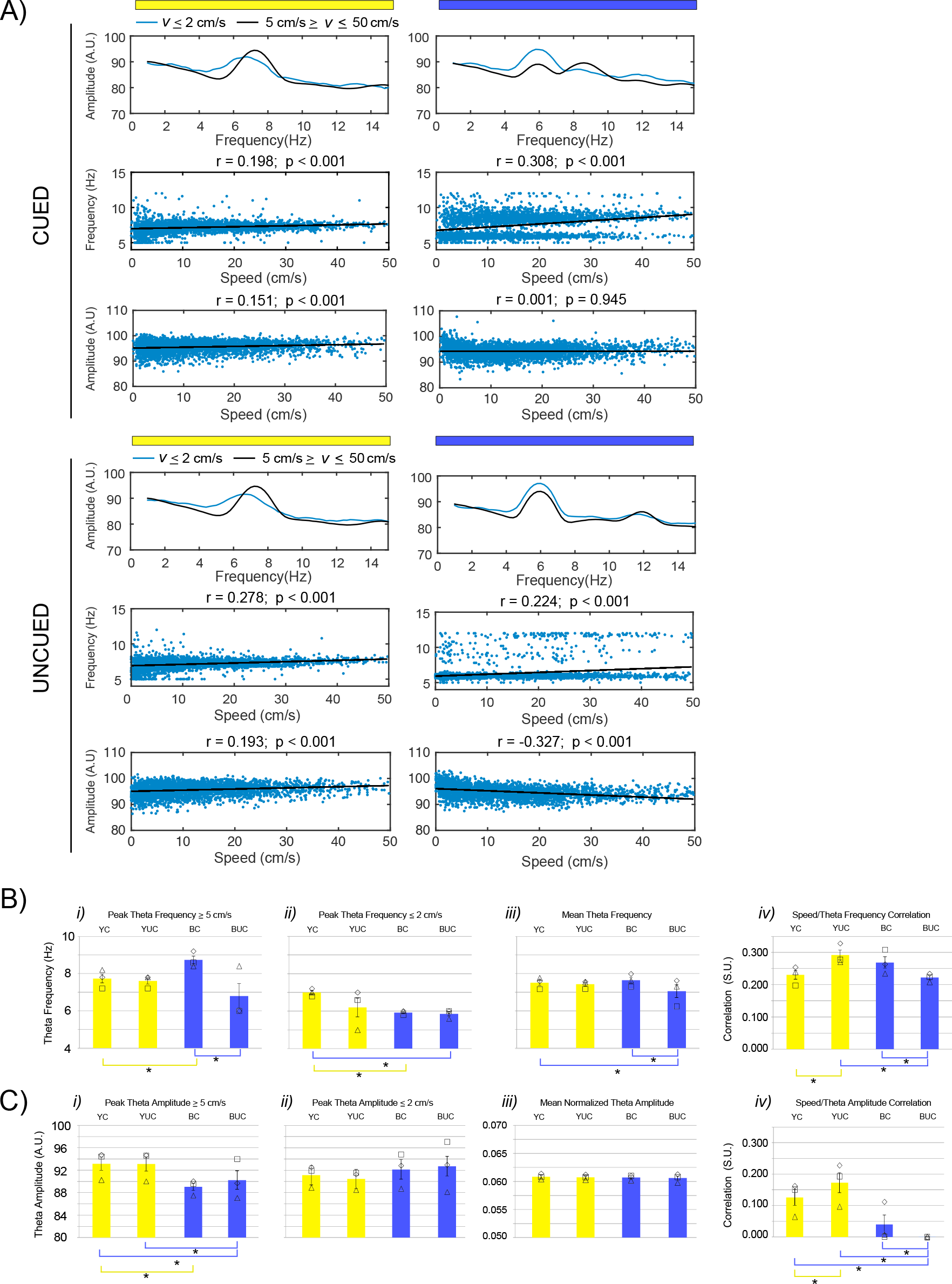
Experiment 2 - Examples of hippocampus speed/theta property dynamics relative to Y and B stimulation during cued (Top) and uncued (Bottom) spatial accuracy conditions. **A)** In each condition mean theta amplitude by frequency during slow (≤ 2cm/s = blue lines) and fast (≥ 5 cm/s = black lines) velocities (*v*) during Y and B MS stimulation is shown. The correlation between movement speed and the frequency and amplitude of each theta cycle (blue dots) for each condition is illustrated while the line of best fit (black line) and the corresponding correlation coefficients and p values are shown above each plot. Linear relationships between theta frequency and amplitude are apparent during Y stimulation in both spatial accuracy conditions, particularly during the uncued condition. In contrast, the speed/theta amplitude relationship is perturbed during B stimulation in both spatial accuracy conditions. The effect of B stimulation has a complex relationship with speed/theta frequency in each task condition. In the cued session, mean theta peaks occur at the MS input frequency at 6 Hz when the rat is moving less than 2 cm/s. At faster speeds, theta toggles to a higher frequency that still increases linearly with speed. This endogenous theta signal is weaker in the uncued session and theta frequency primarily matches the MS input frequency at 6 Hz. **B)** GEE analysis of theta frequency properties across animals (P2 = Triangle, P3 = Square, P4 = Diamond) in relation to stimulation conditions. Yellow and blue lines beneath each plot indicate referential comparison against YC and BUC respectively. Mean peak theta frequency during speeds ≥ 5 cm/s (*i*) was significantly faster during BC than baseline or BUC sessions, indicating competition with endogenous theta. For 2/3 rats the artificial theta signal at 6 Hz was more likely to supersede endogenous theta during BUC. However, at speeds ≤ 2cm/s (*ii*) the peak theta frequency was closer to 6 Hz in both BC and BUC conditions and thus significantly slower than during baseline in YC. Analysis of mean theta frequency without speed filters (*iii*) shows that there is no difference between BC and YC conditions, suggesting that the competition between endogenous and artificial theta during BC serves to maintain theta frequency at its mean value (~ 7.5 Hz). The mean theta frequency is lower during BUC than both BC and YC conditions, again suggesting that entrainment by artificial theta is improved. In agreement with these results, the dynamic relationship between speed and theta frequency correlations (*iv*) is maintained during BC and is on par with YC and YUC conditions. However, r-values are significantly lower during BUC than both BC and BUC conditions indicating that the increased tendency to oscillate at 6 Hz at all speeds perturbs the linear speed/theta frequency relationship. The significant increase in r-value between YC and YUC conditions implies background network changes that occur in relation to increased cognitive demand during uncued spatial accuracy. **C)** GEE analysis of theta amplitude properties across rats in relation to stimulation conditions. Mean peak theta amplitude during speeds ≥ 5 cm/s (*i*) was significantly reduced during BC and BUC conditions in comparison to baseline. Although amplitude was somewhat elevated in these conditions during speeds ≤ 2cm/s (*ii*), they were not significantly different from baseline. While a lack of significant differences was also observed for normalized mean theta amplitude across conditions (*iii*), the tendency for elevated amplitude during slower speeds and decreased amplitude during faster speeds significantly disrupts the speed/theta amplitude relationship (*iv*) during B stimulation.

Without speed filtering, GEE analysis revealed that B stimulation significantly decreased mean peak theta frequency (YC = 7.67 ± 0.196 Hz; YUC = 7.60 ± 0.163 Hz; BC = 6.73 ± 0.599; BUC = 5.93 ± 0.054 Hz; p < 0.0001). Peak theta frequency during BC was not different from baseline conditions during YC (p = 0.54) or BUC (p = 0.221) while BUC closely matched the 6 Hz input frequency and was significantly slower than YC (p < 0.001). B stimulation efficacy was therefore better during the uncued task than the cued task and indicates task condition can significantly influence hippocampal entrainment. However, this result doesn’t account for the influence of animal speed. At speeds ≥ 5 cm/s (Fig. 5A), a significant main effect of stimulation condition was found (p < 0.0001) where mean peak theta frequency during BC was significantly faster than during BUC (p = 0.027) and YC (p = 0.003) (Fig. 5B*i*). This increase in peak theta frequency during BC suggests that accelerated endogenous theta was more likely to dominate the septal input frequency in this condition, but not during the BUC condition. During BUC, 2/3 rats had peak theta frequencies that matched septal input at 6 Hz. For these animals entrainment by artificial theta stimulation was improved during the uncued task, even while moving at speeds ≥ 5 cm/s. For the remaining animal (P2), the endogenous peak theta frequency was still more dominant than the septal input frequency. One difference between P2 and the other rats was that the GFP expression, while the densest of the three animals in the diagonal band of MS, was less present in the horizontal limb of the MS. It is possible then that to fully supersede endogenous LFP theta oscillations at all speeds, ChR2 may also need to be present in both horizontal and vertical limbs of the MS.

At movement speeds ≤ 2 cm/s a significant main effect of condition (Fig. 5B*ii*) was also found for mean peak theta frequency (p < 0.0001). Importantly, peak theta in both BC and BUC conditions matched 6 Hz input frequency for 3/3 animals and were significantly slower than baseline conditions during YC (p < 0.0001 in both cases). Task conditional entrainment of mean peak theta frequency was not found at slow movement speeds when comparing BC and BUC (p = 0.644). As in Experiment 1, this result indicates that entrainment of hippocampal theta oscillations at septal input frequencies is more likely at slow speeds or when the animal is at rest, when there is less competition with endogenous theta.

The end effect of the competition between endogenous and artificial theta, particularly during BC, is to maintain a linear relationship with speed and an average frequency at approximately 7.5 Hz. Further supporting this interpretation, a significant effect of stimulus condition was found on the mean theta frequency in the absence of speed filtering (p = 0.036), where there was no difference between mean theta frequency during YC and BC (p = 0.298) (Fig. 5B*iii*). In keeping with the notion that hippocampal entrainment by artificial theta was improved during the uncued condition, mean theta frequency was significantly slower during the BUC condition than either BC (p = 0.006) or YC conditions (p = 0.042). Similarly, a significant main effect of stimulation condition was found for the linear relationship between speed and theta frequency (p < 0.0001) where the speed/theta frequency relationship during BC was equal to baseline conditions during YC (p = 0.191). In contrast, correlations were significantly reduced during BUC in comparison to both BC (p = 0.001) and YUC (p < 0.0001). This further indicates that the network’s ability to maintain a dynamic relationship with animal speed is perturbed during the BUC session. Yet, this perturbation is not extreme as it was not significantly different from baseline conditions in YC (p = 0.549) (Fig. 5B*iv*). A suggestion as to the origin of entrainment differences between the uncued and cued task variants comes from comparison between the speed and theta frequency correlations during YC and YUC. The significant increase in the correlation value between YC and YUC (p < 0.0001) may reflect network level changes associated with increased cognitive demand [34] during the uncued spatial accuracy condition.

In contrast to a previous report using selective optogenetic stimulation of PV interneurons in transgenic mice [24], pan-neuronal stimulation in this experiment did influence the relationship between animal speed and both theta frequency and amplitude. Without speed filtering, B stimulation had no effect on normalized mean theta amplitude (p = 0.964; Fig. 5C*iii*) or peak theta amplitude (YC = 92.62 ± 1.15 A.U.; YUC = 92.17 ± 1.36; BC = 88.85 ± 0.82; BUC = 90.41 ± 2.02; p = 0.120). However, a significant main effect was found during movement speeds ≥ 5 cm/s (p = 0.004; Fig. 5C*i*). Specifically, mean peak theta amplitude was reduced in both BC (p < 0.0001) and BUC sessions (p = 0.009) in comparison to baseline. In contrast, when movement speeds were ≤ 2cm/s, mean theta amplitude was elevated during both BC and BUC, although these differences were non-significant in relation to baseline (p = 0.79; Fig. 5C*ii*). Given the tendency for stimulation to reduce theta amplitude at fast speeds and elevate the signal amplitude at slow speeds (see Fig. 5A), we used GEE analysis to test for possible perturbation of the linear relationship between speed and theta amplitude. A significant main effect of condition was found for the correlation of speed and theta amplitude (p < 0.0001; Fig. 5C*iv*). Unlike the relationship between speed and theta frequency, in comparison to baseline the positive linear correlation between speed and theta amplitude was significantly reduced during BC (p < 0.0001) and completely abolished during BUC (p < 0.0001). This was largely due to the inversion of the speed/theta amplitude relationship during BUC (see example in Fig. 5A). In this analysis negative correlation values were transformed into small positive values so that gamma with log link models could be fit to the correlation data for GEE analysis.

The MS signals corresponding to the example in Fig. 5A are shown in Fig. 6A. Applying the same approach to theta band properties in the MS, there was a significant main effect of stimulation condition for mean theta frequency without speed filtering (p < 0.0001; Fig. 6B*i*). There was a significant difference between BUC and baseline conditions in YC and YUC (p < 0.0001). As the mean theta frequency was 6 Hz during B stimulation regardless of task, there was no significant difference in mean theta frequency between BC and BUC (p = 0.750). Analyses with and without speed filters demonstrated a clear increase in normalized signal amplitude during B stimulation conditions relative to YC and YUC (p < 0.0001; Fig. 6B*ii*). There was no significant task effect for theta signal amplitude during BC and BUC (p = 0.49). A significant main effect of condition was present with regard to speed/theta frequency (p < 0.0001), where speed/theta correlations were abolished during B stimulation and were significantly lower during BUC than YC (p = 0.001) or YUC (p < 0.0001; Fig. 6B*iii*).

**Figure 6:**
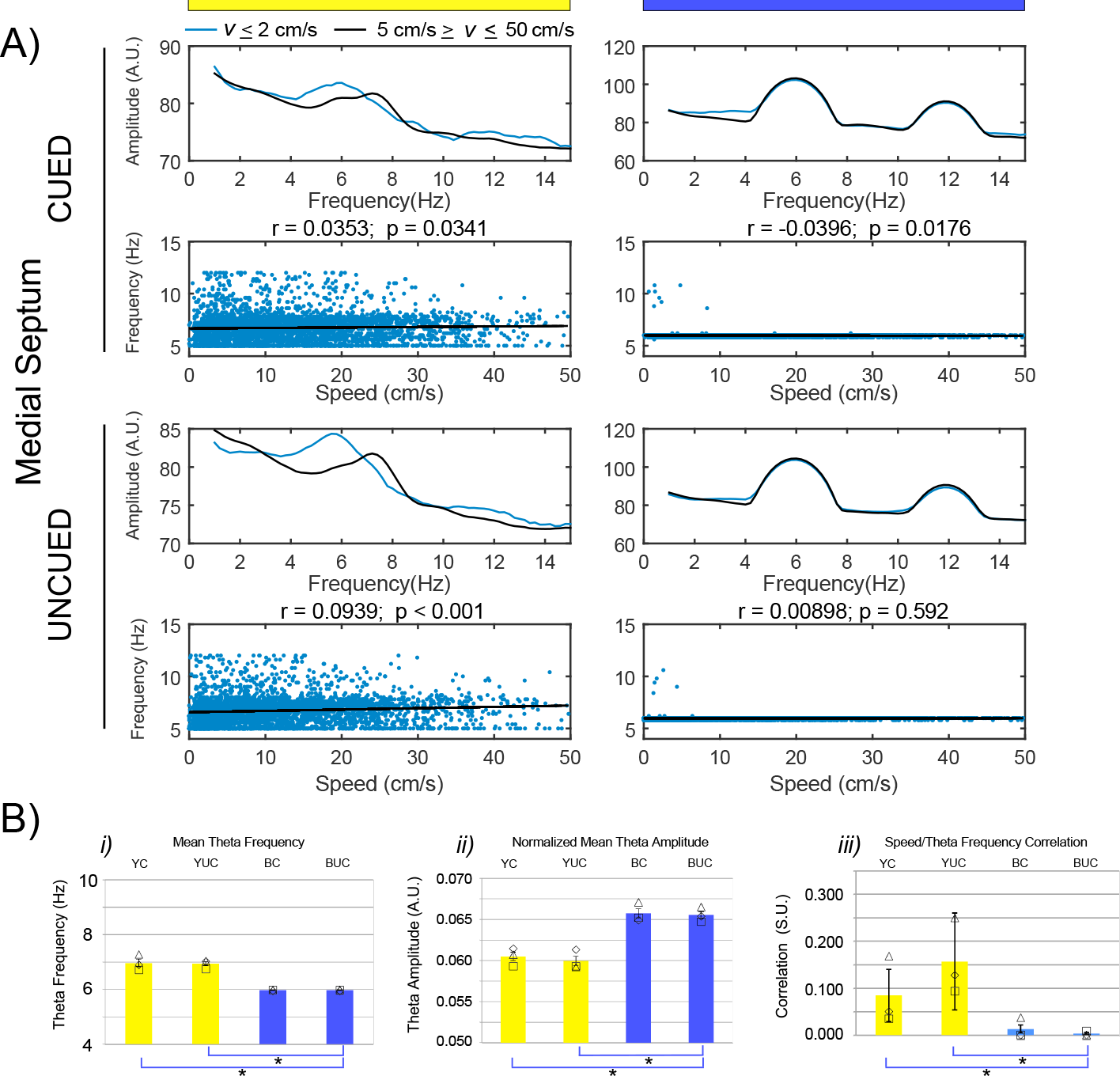
Experiment 2 - As in Fig. 5, example plots of simultaneously recorded MS theta signals. **A)** During Y stimulation speed/theta correlation is less robust in the MS than the hippocampus, although there is a marked increase in r-value during the uncued session. During B stimulation, theta signals are fixed at the 6 Hz input frequency regardless of speed. **B)** Mean hippocampal theta frequency (*i*) and mean normalized theta amplitude (*ii*) without speed filters in each condition for all animals. B MS stimulation reduces the mean frequency to 6 Hz while significantly increasing theta amplitude. The speed/theta frequency relationship is abolished during B stimulation.

While the majority of this report focuses on 6 Hz stimulation of the MS, we also did a limited series of experiments using 10 Hz stimulation in the same animals. The mean and standard errors of the same variables described in the 6 Hz stimulation analyses are reported in Supp. Table 2. In brief, during BC there was a tendency of the endogenous hippocampal theta signal to shift downward from the artificial input at 10 Hz at movement speeds greater than 20 cm/s (Supp. Fig. 6A). This led to an inversion of the speed/theta frequency correlation. As was the case with 6 Hz stimulation experiments, during BUC the relationship between the hippocampal LFP and septal input was more linear and independent of animal speed (Supp. Fig. 6B). However, across animals, differences in task conditional efficacy during 10 Hz entrainment were not statistically significant. Both peak and mean theta frequency during BC and BUC conditions tended to resemble the septal input frequency more than endogenous theta. This implies that network compensation is less likely at septal input frequencies at the faster end of the theta bandwidth.

Finally, hippocampal LFPs were recorded from an additional 3 animals that foraged for pellets in the open field (Supp. Fig. 7). Speed/theta frequency correlations were on par with the YC condition where recordings were carried out in the absence of cognitive demand (Supp. Table 3). In addition, there were no significant differences in speed/theta frequency or other theta property variables between 2 different exposures to the open field. This speed/theta frequency stability implies that the significant increase in correlation values between YC and YUC in the previous experiment was not an effect of re-introduction to the arena and was more likely a consequence of increased cognitive demand.

We have therefore shown that artificial theta oscillations generated through MS optogenetic stimulation supersedes endogenous hippocampal oscillations when the animals are moving slowly or at rest in both experiments 1 and 2. Entrainment of hippocampal theta oscillations during faster movements is more complex and influenced by task. This strongly suggests altered microcircuit dynamics in the septo-hippocampal circuit during performance of the uncued rather than the cued spatial accuracy task. We have also sown that MS stimulation disrupts the dynamic relationship between animal speed and theta amplitude.

### MS stimulation perturbs endogenous theta phase preference

As septo-hippocampal LFP entrainment efficacy was affected by task conditions, we aimed to determine if cognitive demand also influenced temporal processing of CA1 pyramidal cells relative to artificial or endogenous theta oscillations in the hippocampus and MS. In all 4 simulation/task conditions we calculated the mean angle and magnitude of phase preference relative to hippocampus and MS theta as determined by the resultant vector lengths of cells pooled across animals (n =166; Fig. 7A-C) as well as for cell population ensembles from each of the 3 animals (n = 22, 57, and 87; Fig. 7D-E). Illustrations of circular statistics for phase preference analysis of pooled cell data relative to theta bandwidth signals in the hippocampus and MS during each stimulation condition are shown in Fig. 7A and B respectively. A table showing Watson-Williams multi-sample test comparisons across conditions can be found in Fig. 7C. Pyramidal cells demonstrated a significant phase vector at the ascending phase of theta during both YC and YUC. While the phase angle did not significantly differ between these 2 conditions, it is notable that the population vector (RBAR) during YUC almost doubled in comparison to YC. Endogenous theta phase preference was non-significant during both BC and BUC sessions and significantly differed from their corresponding baseline stimulation sessions during YC and YUC.

**Figure 7:**
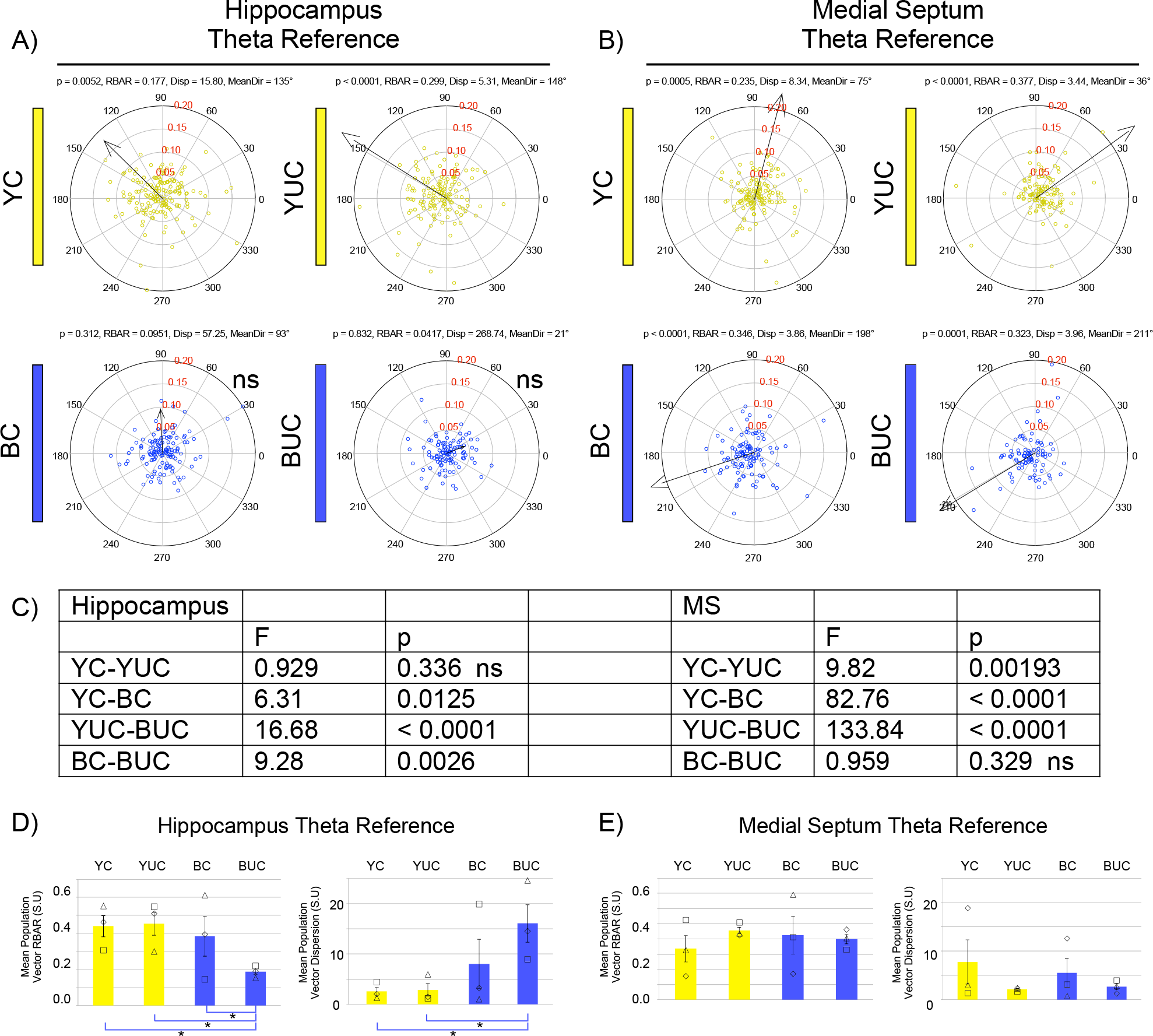
Experiment 2 - Circular statistics for hippocampal and MS theta phase preference of CA1 pyramidal cells per stimulation condition (YC = 6 Hz Yellow stimulation during cued spatial accuracy; YUC = 6 Hz Yellow stimulation during uncued spatial accuracy; BC = 6 Hz Blue stimulation during cued spatial accuracy; BUC = 6 Hz Blue stimulation during uncued spatial accuracy). Each circle represents the angle and magnitude of resultant theta phase vector (RBAR) for an individual pyramidal cell. The arrow indicates the angle and magnitude of the resultant theta phase vector for the population. Discontinuous arrowheads indicate resultant vectors > 0.2. RBAR levels for the circular plot are shown in red. The p-values, population RBAR values, levels of population dispersion and mean vector angle (MeanDir) are shown above each plot in **A-B**. Non-significant vectors (ns) are noted. **A)** Between YC and YUC, the hippocampal theta phase vector is stable although the RBAR increases and the population dispersion decreases. Phase preference is significantly perturbed during BC and abolished during BUC. **B)** Between YC and YUC the RBAR of the septal phase vector increases and the phase angle shifts closer to the trough at 0°. In contrast to the hippocampal theta vector, MS theta shifts approximately 180° during 6 Hz B stimulation. This difference is particularly striking between the YUC and BUC sessions. **C)** Supp. Table 3 shows the results of Watson-Williams circular tests comparing YC-YUC, YC-BC, YUC-BUC and BC-BUC relative to each reference signal. **D-E)** Mean and standard error of the population vector RBAR (Left) and population dispersion (Right) for all 3 rats relative to hippocampal theta and septal theta. **D)** Hippocampal theta RBAR and dispersion levels were variable across rats during BC but were universally degraded during BUC. **E)** No significant effects of stimulation condition were found relative to septal theta. Again, there was less variability across animals during uncued spatial accuracy for both RBAR and dispersion measures.

With regard to septal theta, there was a significant difference between YC and YUC conditions where the population phase vector shifted back approximately 40 degrees, toward the theta trough. In contrast to hippocampal theta, a significant population vector was maintained during 6 Hz B stimulation in both the BC and BUC sessions. However, these vectors were flipped toward the descending phase of theta representing a striking ~ 180° phase shift between YUC and BUC sessions. While significant phase differences were found between Y and B stimulation during the cued and uncued sessions, phase vectors were stable between BC and BUC sessions.

GEE analysis of the length of the resultant phase vectors (RBAR) for the cell populations referenced to hippocampal theta in each animal (Fig. 7D Left) found a significant effect of stimulation condition (p < 0.0001). There was no change in RBAR between YC and YUC sessions (p = 0.194), yet a significant reduction was identified when comparing both YC and YUC sessions to the BUC session (p < 0.0001). RBAR during BC was variable between animals and, while only marginally different from BUC (p = 0.053), was also similar to YC (p = 0.379). A complementary analysis of population level dispersion (Fig. 7D Right) also found a significant main effect of stimulation condition (p < 0.0001). Again, while YC and YUC had similarly low levels of dispersion, there was a significant increase in dispersion levels during BUC (p < 0.0001) in comparison to both YC and YUC sessions.

When the same cells are referenced to MS theta and compared across animals, there was no significant main effect of stimulation condition (Fig. 7E) on either the length of the resultant phase vector (p = 0.717) or levels of dispersion (p = 0.101). Taken together with the circular statistics in Fig. 7A-B, this result further indicates that the temporal organization of hippocampal cells by endogenous theta phase is superseded by the artificial septal stimulation, albeit at a significantly different phase angle.

As a further exploration of the influence of cognitive demand on phase preference, we recorded cells (N = 158) during a foraging task with 3 additional animals (n = 62, 29 and 67) and analyzed spike activity relative to local hippocampal theta oscillations (Supp. Fig. 8A-B). Circular statistics from the pooled cells revealed a significant resultant population vector (RBAR = 0.169, Disp = 15.46, p = 0.01) with a mean direction of 213°. Watson-Williams tests revealed significant differences between theta phase preference during foraging and both the cued (F = 30.83, p < 0.0001) and uncued (F = 23.73, p < 0.0001) versions of the spatial accuracy task. While the dominant phase angle is different between YC and foraging, RBAR and dispersion levels are similar (RBAR = 0.177, Disp = 15.8). The increased RBAR and lower dispersion levels during YUC (RBAR = 0.299, Disp = 5.31) may therefore be indicative of increased temporal coordination within CA1 in response to increased cognitive demand with regard to spatial processing.

While a reduction in CA1 theta phase preference was variable during BC, temporal organization relative to local oscillations was universally perturbed across rats during BUC. In this stimulation condition hippocampal theta phase preference was superseded by artificial septal theta oscillations, but shifted 180° to the opposite phase of septal theta preference during YUC. These results therefore imply a temporally ordered hierarchy among hippocampal projections that are governed by theta phase and corresponding cognitive demands. Hippocampal pyramidal cells are more attuned to artificial MS stimulation than endogenous hippocampal theta oscillations, particularly during the BUC condition.

### MS entrainment of pyramidal cell burst-timing mirrors task effects on LFP entrainment

We applied the same GEE methodology to the results of autocorrelation analyses to test for possible effects of stimulation condition on the organization of timing intervals between spike trains from CA1 pyramidal cells (n = 144) from 3 animals (n = 23, 64 and 57). Examples from an individual cell during each stimulation condition with corresponding autocorrelation histograms and modulation depth frequency from the ACG analysis are shown in Fig. 8A. As this analysis was sensitive to rate changes, we first analyzed for any main effect of mean overall rate by stimulation conditions. Although there was a decrease in mean overall rate during YUC, rate was not significantly different between baseline conditions in comparison to B stimulation during cued sessions or uncued sessions. Stimulation did not alter overall mean firing rate.

**Figure 8:**
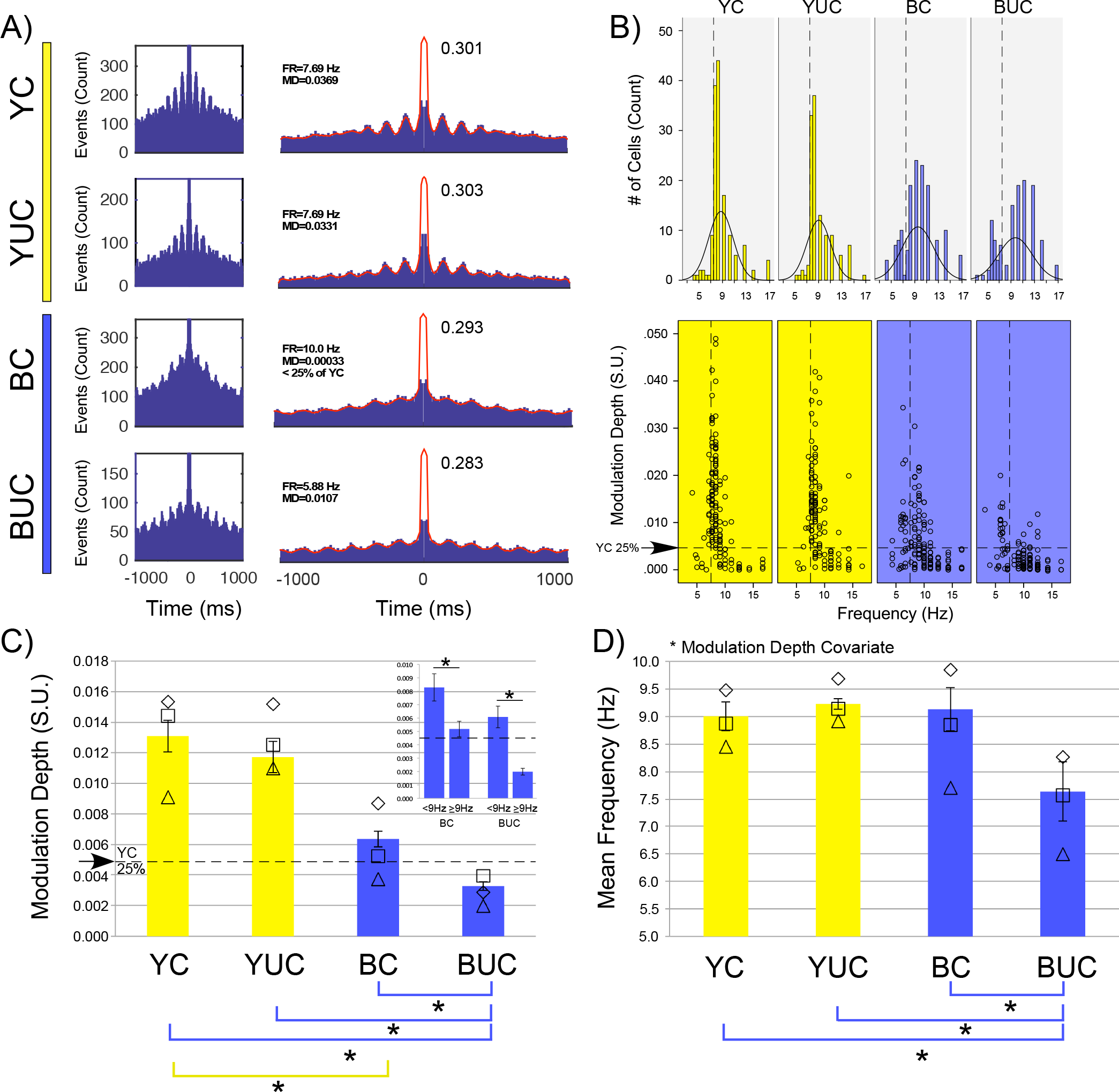
Experiment 2 - Stimulation and task conditional effects on the timing of CA1 pyramidal neuron inter-spike intervals. **A)** Example autocorrelation histograms (Left) and autocorrelograms (Right) for a pyramidal cell during 6 Hz Y MS stimulation in the cued (YC) and uncued (YUC) tasks and during 6 Hz B MS stimulation during the cued (BC) and uncued (BUC) tasks. Frequency (FR), modulation depth (MD), and peak autocorrelation (right of each peak) are shown in each condition. Periodicity and modulation depth for the cell are constant during both baseline conditions while the cell is poorly modulated during BC. During BUC there were prominent changes in modulation depth and periodicity indicating entrainment at the septal input frequency of 6 Hz. **Top B)** Histogram representing the number of cells by modulation frequency in each task and stimulation condition with overlaid normal distribution curves. The dashed line serves as a reference point for the mean LFP theta frequency during YC, YUC and BC at 7.5 Hz. The majority of cells during baseline conditions are clustered ~ 1 Hz ahead of the mean theta frequency. While task condition has little effect on the distribution of cell periodicity during YC and YUC, during BC the distribution shifts to higher frequencies and has a peak at 9 Hz. During BUC the distribution appears bimodal with 2 populations separated at approximately 7.5 Hz and a peak at 11 Hz. **Bottom B)** As modulation frequency can be overestimated with low modulation, the frequency data from D are presented in a scatterplot against modulation depth for each condition. Each circle represents data from a single pyramidal cell and the dashed line in the Y-axis again serves as a reference for mean LFP theta during YC, YUC and BC at 7.5 Hz. The dashed line in the X-axis denotes the modulation depth cut-off for the bottom quartile of cells during YC baseline (Tukey’s hinge = 0.0045). A separation of cells into 2 populations exhibiting modulation frequencies either > or < 7.5 Hz is evident during BC. During BC 30% of the cells shift toward the input frequency at 6 Hz while the remainder of well-modulated cells shift toward faster frequencies or exhibit attenuated modulation. During BUC, the subset of cells < 7.5 Hz are still well modulated while those > 7.5 Hz are not. Spike timing results therefore reiterate the competition between endogenous and artificial theta frequency during BC as well as the attenuation of the influence of endogenous theta during BUC. **C)** Mean and standard error of inter-spike interval modulation depth across animals during each stimulation and task condition. The dashed line in the X-axis again denotes the lower quartile for modulation depth during YC. Mean modulation depth decreases significantly during B stimulation conditions. Modulation depth during BUC is significantly lower than all other conditions and the mean depth for each animal is lower than the bottom quartile in the YC baseline session. Analysis of variance for the modulation depth of all cells that were < or ≥ 9 Hz (Inset) reveals a separation of frequency entrainment by condition. In both BC and BUC approximately 70% of all cells exhibit beat frequency ≥ 9 Hz and are significantly less modulated than cells at beat frequencies ≤ 9 Hz. Only cells that modulated at ≥ 9 Hz tend to have a negligible modulation depth during BUC. **D)** Mean and standard error of modulation frequency with modulation depth as a covariate during each stimulation and task condition. In both Y baseline conditions and BC the mean frequency is approximately 9Hz but decreases significantly during BUC.

A histogram showing the distribution of frequencies for pooled cells in each stimulation condition is shown in Fig. 8B (Top). The inter-spike interval periodicity during both baseline conditions for the majority of the cells was ~ 1 Hz ahead of the mean theta frequency. This phenomenon was explored in detail in a recent study [24]. The frequency distribution is unaffected by task demands in the baseline sessions yet was dramatically altered during B stimulation. Frequencies were more normally distributed within theta band during BC where most of the cells were at higher burst frequencies. During BUC the distribution separated at approximately 9 Hz and was more bimodal. Modulation frequency measurements also exhibited a significant interaction of stimulation condition with modulation depth (p < 0.0001), where all sessions were significantly different from BUC (p < 0.01). In addition, as noted previously, frequency measures tended to be overestimated when modulation depths were low. We therefore represented the dataset as a scatterplot comparing modulation depth against frequency (Bottom Fig. 8B). The frequency distribution was referenced against the mean LFP theta frequency across animals during the YC baseline session. Likewise, the distribution of the modulation depth was referenced to the 25^th^ percentile mark during YC (Tukey’s hinge = 0.0045, see boxplots for YC and YUC sessions in Supp. Fig. 9A-B). Mirroring the LFP result, burst-timing frequencies during BC were split between the input frequency at ~ 6 Hz and frequencies faster than the mean LFP frequency. Remarkably, during BUC the majority of cells fell below the 25^th^ percentile mark. The cells that remained significantly modulated exhibited frequency values close to septal input at 6 Hz.

GEE analysis revealed a significant main effect of pyramidal cell modulation depth (Fig. 8C) by stimulation condition (p < 0.0001). The modulation depth was lowest during BUC and significantly different from all other sessions (p < 0.0001). BC was also significantly different from YC (p < 0.0001). Using modulation depth as a covariate, GEE factorial analysis found a significant effect of stimulation condition across animals (p = 0.007, Fig. 8D) where BUC was significantly reduced in comparison to YC (p = 0.023), YUC (p = 0.04) and BC (p = 0.01). This result is similar to stimulation effects on mean theta frequency (Fig. 5B) where the efficacy of the artificial stimulation is improved during BUC.

In response to this observation, we conducted an additional test for modulation depth for all cells during BC and BUC. Cells were separated into 2 populations by modulation frequency, either ≥ 9 Hz or < 9 Hz (Inset Fig. 8C). We chose this frequency based on the results of Fig. 8B-C, where the 2 populations appear to separate. During BC, cells with a modulation frequency < 9Hz (38%, Mean = 0.00828 ± 0.0010) were compared to cells with a modulation frequency > 9 Hz (62%; Mean = 0.00518 ± 0.00058) by one-way analysis of variance. Cells with a modulation frequency < 9 Hz were found to be significantly more modulated than those ≥ 9 Hz (p = 0.005). Although all cells were less modulated during BUC, we found a similar modulation pattern as in BC. An additional one-way analysis of variance found that cells with a modulation frequency < 9Hz (31%, Mean = 0.00607 ± 0.0008) were again significantly more modulated (p < 0.0001) than cells with a modulation frequency > 9 Hz (69%; Mean = 0.00199 ± 0.00025). During BUC then, cells that remained robustly modulated were more likely to exhibit beat frequencies matching septal input.

As in Experiment 1, cells with modulation frequencies either ≥ 9 Hz or < 9 Hz during BC were traced back to the YC session to determine if their baseline mean firing rates could predict response to B stimulation. Unlike the results during the flowerpot recordings, mean firing rate did not predict stimulation responsive cells (1.21 ± 0.144 Hz vs. 1.29 ± 0.13 Hz; p = 0.719).

We have shown a close correspondence between both the burst-timing properties of hippocampal neurons and synaptic oscillations within the theta bandwidth. At both cell and LFP levels, the efficacy of entrainment at the septal input frequency was significantly affected by stimulation condition and task. The same proportion of cells in the cued and uncued sessions responded robustly to B stimulation. Unlike experiment 1, these cells were not more active than other cells during baseline, suggesting a more complex interaction with active navigation.

### Stimulation induced changes to LFPs and the temporal organization of spike-timing do not alter place cell firing fields

As we found significant stimulation induced alterations in the temporal organization of hippocampal pyramidal cells, we asked if these changes influenced the spatial firing of place cell firing fields. A total of 103 cells met criteria for place cell analyses from 3 rats (n = 18, 44 and 41). Examples of several simultaneously recorded place cell firing fields across stimulation conditions are shown in Fig. 9A. As summarized in Fig. 9B, GEE analysis found no significant main effect of stimulation conditions for mean field coherence, mean overall firing rate, mean firing field stability, mean field size, mean in-field firing rate, and mean out-field firing rate. We therefore conclude that perturbation of hippocampal theta oscillations through septal manipulation do not interfere with the spatial firing of hippocampal place cells, in accordance with previous studies that used MS or systemic pharmacological interventions to perturb hippocampal theta dynamics [21, 49]. Although the temporal organization of cell activity has been altered by septal stimulation, the spatial code remains intact.

**Figure 9:**
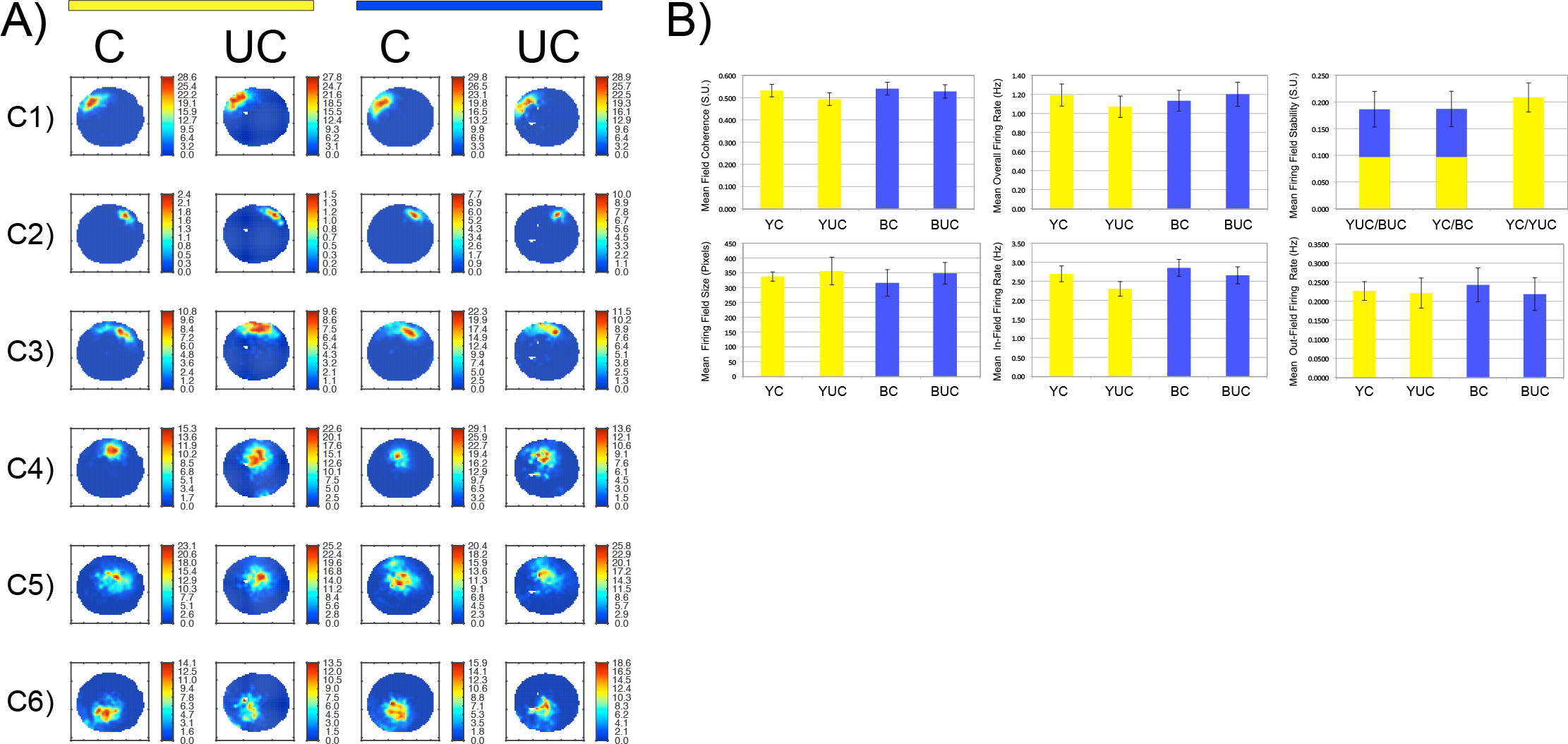
Experiment 2 - **A)** Examples of simultaneously recorded place cell firing fields during each task and stimulation condition. **B)** Results of field property analysis between conditions. Although LFP, phase preference and the temporal organization of spike timing are significantly altered by B stimulation and task conditions, no significant changes were found with regard to place cell firing field properties (mean field coherence, mean overall rate, mean firing field stability, mean firing field size, mean in-field firing rate or mean out-field firing rate). Although temporal coordination is altered, spatial coding remains the same.

### Superseding endogenous theta oscillations attenuates spatial accuracy in the uncued task condition

We have shown that MS stimulation affects the endogenous LFP theta band, theta phase preference and the inter-spike interval of pyramidal cell action potentials, but spares the spatial tuning of hippocampal place cells. While the spatial code in the hippocampus is maintained, the temporal organization of cell activity is predominantly driven by an artificial theta signal. We therefore ask if the temporal organization of hippocampal cells relative to local, endogenous theta oscillations is necessary for self-localization relative to a goal zone in the uncued spatial accuracy task. For each of the 3 animals, the series of stimulation conditions (YC, YUC, BC and BUC) was repeated 3-4 times on different days. Examples of behavioral performance via speed and dwell-time in each stimulation condition are shown in Supp. Fig. 10A-C. Across animals, key variables that illustrated the effect of 6 Hz stimulation on accuracy performance were percentage of time spent in each arena quadrant, search accuracy in the target quadrant, number of rewards, number of entrances, and the ratio of rewards to goal zone entrances. For each variable, data are demonstrated for each experimental series per individual rat in Fig. 10 (A,C,E,G,I) as well as overall mean and standard error in Fig. 10 (B,D,F,H,J).

#### Quadrant dwell time

Across animals B stimulation generally had no significant impact on the distribution of time spent in each quadrant (Fig. 10A). There was no main effect of stimulation condition (Fig. 10B), in the Target (T; p = 0.483), Counterclockwise (CCW; p = 0.12), or Clockwise (CW; p = 0.703) quadrants. While GEE found a significant main effect of condition in the quadrant opposite the target quadrant (O; p = 0.016), this was driven solely by the significant difference between the BC and BUC sessions (p = 0.025). Due to the high number of rewards in the BC session (see below), this result is likely a reflection of increased foraging in the opposite quadrant in comparison to the BUC session.

#### Search accuracy in target quadrant

As described above, the distance of slow speed movements from the goal zone center were used as a measure of search accuracy within the target quadrant. B stimulation during the uncued sessions appeared to increase the distance of goal searching behavior from the center of the goal zone in P4 and P2 but not P3 (Fig. 10C). GEE analysis across rats (Fig. 10D) revealed a significant main effect of stimulation condition (p < 0.0001) where stimulation during BUC increased the distance of search behavior from the goal center relative to all other conditions (YC = 18.05 pixels ± 1.19, p < 0.0001; YUC = 21.08 pixels ± 1.90, p = 0.046; BC = 18.64 pixels ± 1.48, p < 0.0001; BUC = 26.73 pixels ± 1.86). This result suggests that in the uncued task B stimulation leads to goal zone searches that are approximately 9 pixels (~ 3 cm) further from the target center than during the YUC session.

**Figure 10:**
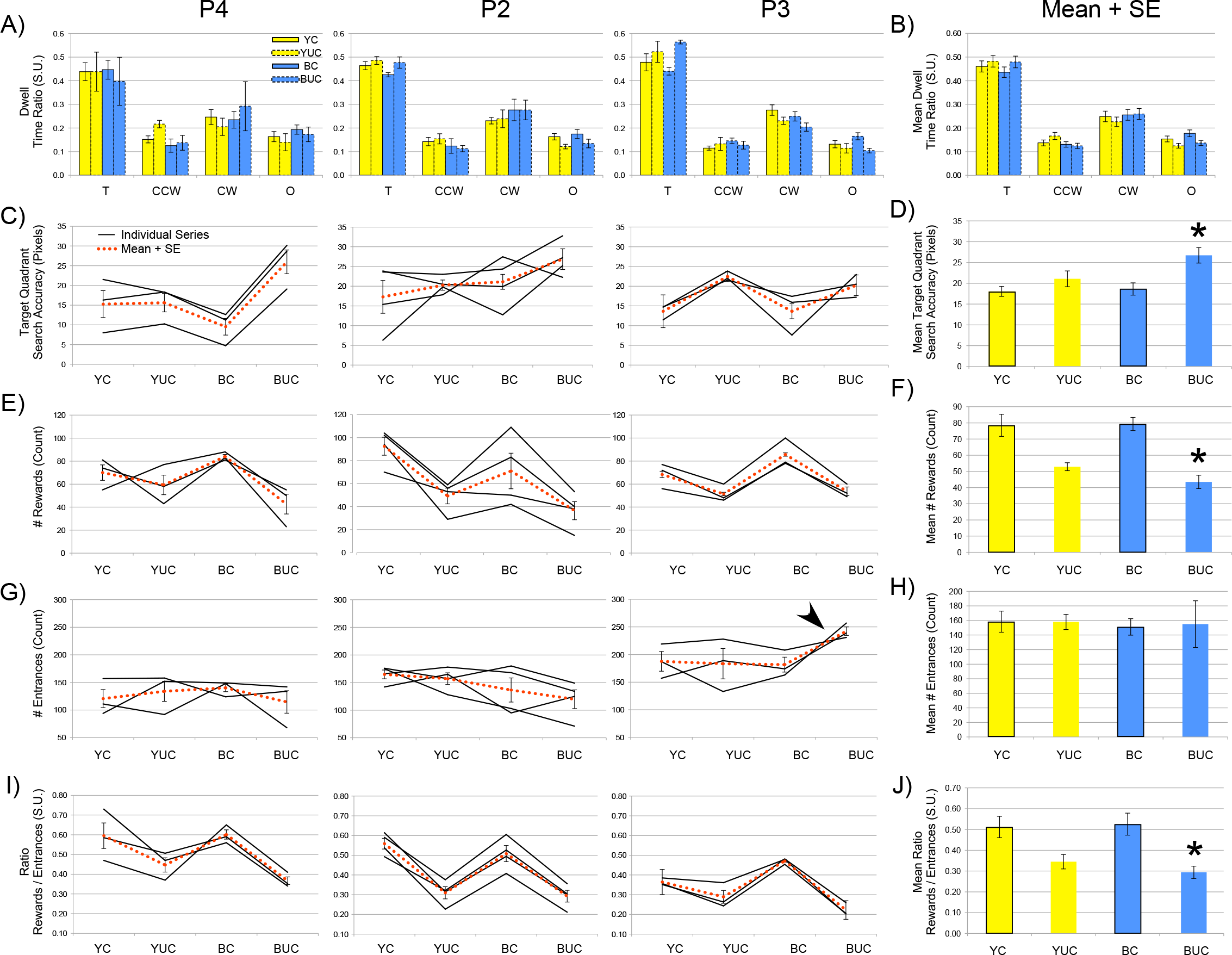
Experiment 2 - Spatial accuracy behavior per rat and GEE analysis across rats. **A)** Mean dwell-time ratio per quadrant for rats P4, P2 and P3. B) Mean and SE of dwell-time ratio for all sessions across rats reveals that all rats spent a similar amount of time in the goal zone quadrant regardless of stimulation condition. **C,E,G,I)** Spatial accuracy performance measures for individual series as well as mean and SE across series per individual rat of Spatial accuracy (Target quadrant search accuracy, Number of rewards, Number of goal zone entrances, Ratio of rewards to entrances). Unlike the other 2 rats P3 did not exhibit a change in search accuracy and the number of rewards in comparison to baseline sessions. However, this rat did make significantly more entrances into the goal zone during the BUC session (Arrow) denoting uncertainty regarding the goal zone location as well as a different strategy than the other 2 rats. **D,F,H,J)** Mean and SE for each behavioral measure across rats. During the BUC condition, GEE reveals a significant decrease in search accuracy, a significant decrease in the number of rewards and a significant decrease in the ratio of rewards to entrances. Artificial entrainment of the septo-hippocampal circuit and the perturbation of the temporal coordination of hippocampal neurons causes the approximation of the goal zone location to be less precise but spares the ability to navigate to the correct quadrant.

#### Number of rewards

As with search accuracy, B stimulation during the uncued session significantly impacted the number of successfully triggered rewards for P4 and P2, but not P3 (Fig. 10E). Nonetheless, GEE analysis across rats (Fig. 10F) revealed a significant main effect of stimulation condition (p = 0.003) where B stimulation during the uncued task significantly decreased reward number in comparison to all other sessions (YC = 78.5 ± 6.87, p = 0.001; YUC= 52.9 ± 2.46, p = 0.043; BC = 79.3 ± 4.08, p < 0.0001; BUC = 43.5 ± 4.14).

#### Number of entrances

Although search accuracy and number of rewards for P3 were not affected by B stimulation in the uncued task, analysis of the number of entrances revealed that P3 made more entrances during the BUC session (Fig. 10G). This trend was not found for the other 2 animals and GEE analysis did not find a significant main effect of stimulation condition (Fig. 10H; p = 0.989). However, these results suggest that P3 was affected by B stimulation during the uncued task, albeit not in the same manner as P4 and P2. This suggests that P3 had an alternative strategy for triggering rewards during the BUC session that involved getting close to the goal zone, moving its head and the LED around the estimated goal position until the pellet feeder was triggered.

#### Reward/Entrance ratio

The ratio of the number of rewards to the number of entrances further suggests that P3, as well as the other animals, tended to make more entrances for fewer rewards during the BUC session (Fig. 10I). GEE analysis showed that this difference was significant across rats (p < 0.0001) where the ratio was lower during the BUC session than all other sessions (YC = 0.512 ± 0.0516, p < 0.0001; YUC= 0.345 ± 0.0348, p = 0.045; BC = 0.526 ± 0.0530, p < 0.0001; BUC = 0.294 ± 0.0296).

#### Lack of order effect

GEE was also used to compare the potential order effect for the BUC session, for whether it preceded or followed the BC session. No significant order effect was found for the percentage of time spent in the Target (p = 0.510), Clockwise (p = 0.527), Counterclockwise (p = 0.689), or the Opposite quadrant (p = 0.456). Likewise, no significant order effect was found for search accuracy (0.614), number of rewards (p = 0.939), or the ratio of number of rewards to number of entrances (p = 0.371).

We conclude that B stimulation during the uncued task did not impair the rat’s ability to find the target quadrant but did attenuate the accuracy with which the rats localized the goal zone. While this effect was subtle, it does suggest that temporal coordination of hippocampal neurons relative to endogenous theta oscillations is not necessary for navigating to the target quadrant but is necessary for precise self-localization within approximately 6 cm of the goal zone center.

## Discussion

We investigated how behavioral state and cognitive demands placed on the septo-hippocampal circuit influenced the entrainment efficacy of both hippocampal local field potentials and single cell activity via nonselective optogenetic stimulation of the MS in wild-type rats. We accomplished this through electrophysiological recordings while titrating behavioral state, when the rats were at rest or actively navigating, as well as cognitive demand in the septo-hippocampal circuit by having the rats perform a cued or uncued spatial accuracy task. Entrainment of local field potentials and cell activity at the 6 Hz septal input frequency was more likely to supplant endogenous theta oscillations when the animal was at rest or moving slowly in either the cued or uncued spatial accuracy task. Generally, it is easier to entrain hippocampal oscillations in the absence of a strong endogenous theta signal. This finding is similar to a previous report [27] where 6 Hz MS stimulation during movements ≥ 2 cm/s revealed complex interactions between the septal pacemaker and local theta inputs. Our experiment found that these interactions were also significantly influenced by the version of the spatial accuracy task.

During both cued and uncued tasks, hippocampal LFP frequencies matched septal input frequency at 6 Hz for all rats when moving at speeds ≤ 2 cm/s. During faster movements in the cued task, the LFP in the theta bandwidth was more likely to ‘toggle’ between 6 Hz input and an endogenous theta frequency at ~ 9 Hz. This apparent competition between artificial and endogenous theta was not only represented in the synaptic current underpinning the LFP but was also found to separate the burst-timing of pyramidal cell action potentials into two separate populations. A simple interpretation of this phenomenon is that displacing theta frequency 1.5 Hz down from the baseline mean theta frequency of 7.5 Hz results in an endogenous theta frequency 1.5 Hz greater than the baseline mean at ~ 9 Hz. Remarkably, although the lower endogenous theta limit was shifted 1.5 Hz upwards from baseline, the endogenous signal still increased linearly with speed. This endogenous signal may be compensatory as it ultimately served to maintain a linear relationship between theta frequency and animal speed. The 2 populations of pyramidal cells separated by burst-timing frequency during the cued accuracy session may therefore be more responsive to either the endogenous or artificial theta signal. In contrast, during faster movements in the uncued task, hippocampal LFP frequencies matching septal input at 6 Hz were more likely to dominate the LFP bandwidth. The LFP result across animals was also mirrored in the burst-timing of pyramidal cell action potentials. Only cells that exhibited a burst frequency near the septal input frequency remained robustly modulated while modulation was attenuated for the remaining majority of cells. In both Experiments 1 and Experiment 2, and in a previous report using optogenetic MS stimulation [24], only 25-30% of the active cells were significantly modulated within 1 Hz of the stimulation frequency. This burst-timing result, as well as the differences in theta phase preference in each stimulation condition, indicates that the active cell population may have been more attuned to septal and subcortical input during the uncued task than the cued version of the accuracy task.

How did the behavioral state and task version influence septo-hippocampal entrainment efficacy? The effect of movement is the easier finding to address and is most likely attributed to the lack of competition between weaker endogenous theta and the artificial theta input. The task effect, competition during the cued task and a more harmonious relationship between endogenous and artificial signals during the uncued task, is harder to address. However, there is a similar precedent. Using a similar goal zone oriented spatial task, Fenton et. al. [50] found that the variance in the firing rate of hippocampal place cells, a phenomenon referred to overdispersion, was significantly reduced during increased spatial demands. This reliability in the firing rate was hypothesized to be due to increased attention to a subset of spatial cues that were relevant for task performance. Likewise, in our experiment increased attention in the uncued version of the task may have reduced the variability in the hippocampal inputs and background processes underlying the active search for the goal zone. This may also explain the increased fidelity between animal speed and theta frequency between the YC and YUC conditions. In addition, during increased cognitive demand OLM interneurons may be more engaged through elevated acetylcholine from septal projections [51]. This could increase inhibitory tone at the cortical inputs into the hippocampus and make the relationship between septal input and local synaptic oscillations more linear. Another similarity between the current and Fenton et al. studies is that the overdispersion phenomenon seen during lower levels of cognitive demand and the ‘toggling’ effect between endogenous and artificial theta both appeared to occur on a seconds timescale.

Stimulation frequency, as well as task condition, may be an important variable to consider with regard to entrainment efficacy. As with 6 Hz stimulation, task influenced the balance of endogenous and artificial hippocampal theta during 10 Hz stimulation. During the cued task peak theta frequency was found to be closer to the mean of baseline theta while during the uncued task the theta peak more faithfully reflected septal input, regardless of speed. However, we did not find a significant task effect on the efficacy of 10 Hz entrainment. Across rats, peak frequency was closer to the input frequency in both cued and uncued sessions. If the competition between endogenous and artificial theta seen during 6 Hz stimulation was indeed a result of circuit compensation for being pushed below theta mean, then it is possible that there are limits to this compensation if the input frequency is further from 7.5 Hz.

Optogenetic stimulation of parvalbumin positive interneurons in the MS of transgenic mice was recently found to pace hippocampal theta frequencies at 10 or 12 Hz stimulation frequencies [24]. While the burst-timing results in this study were similar to our findings, the study did not find changes with regard to phase preference or the resultant length of the phase vector. While the cells no longer showed a consistent relationship with endogenous theta frequency, as determined by the attenuation of the slope of place cell phase precession or with the relationship between animal speed and theta frequency, they found that the relationship between speed and theta amplitude was spared. At both septal input frequencies, we found strong evidence for modulation perturbation in most active pyramidal cells, the disruption of the linear relationship between speed and LFP theta properties, and the abolishment of the theta phase relationship of pyramidal cells during the uncued task. The differences between our study and the Zutshi et al. study [24], particularly with reference to theta phase preference and the linear relationship between animal speed and theta amplitude, was very likely due pan-neuronal septal transduction of ChR2 and the inclusion of cholinergic septal neurons [52–54] rather than just PV interneurons. One advantage to disrupting hippocampal temporal organization, at the level of both the cell and the network, is that it allowed us to ask if these phenomena are necessary for performance of the uncued spatial accuracy task [21–23, 55, 56], or if a stable spatial code is sufficient to guide spatial behavior.

Although rats generally had no difficulty navigating to the target quadrant, stimulation during the uncued task clearly attenuated accurate localization of the goal zone. Relative to baseline in the YUC session, accuracy was off by ~ 2 cm. Importantly, no behavioral effect was found during the cued version of the task. The stimulation therefore had no off-target effects in relation to motor movement or sensory processing. Although 1 of the rats did not exhibit a change in the degree of accuracy or a decrease in the number of rewards, there was a significant increase in the number of entrances. This particular rat used a different strategy in the BUC session that involved sweeping the headstage mounted LED in and out of the goal region until a reward was triggered. This suggests that this rat’s spatial accuracy was also affected but in a different way from the other two rats. As a result, across rats there was a significant decrease in the ratio of the number of rewards to the number of entrances.

The overall attenuation in spatial accuracy across rats was subtle. This result is somewhat surprising given the significant alterations found for multiple levels of temporal coordination in the electrophysiological signatures. One interpretation is that a stable spatial code is sufficient for finding the correct arena quadrant but that temporal organization by dynamic endogenous theta may be necessary for more accurate self-localization within approximately 6 cm. This interpretation of the data is supported by previous experiments that lesioned the medial entorhinal cortex or inactivated the MS and impaired the ability of the rats to estimate linear distances based on self-movement information [19]. Likewise, lesioning of the MS [57] was shown to significantly interfere with the performance of a previously learned spatial task on an elevated circular track. Animals were able to at least recognize a goal location if they stumbled upon it, but otherwise had difficulty in determining goal location. Theta oscillations paced by the MS have therefore been deemed necessary for path integration and determining position in space. In light of our results, endogenous theta oscillations may provide both temporal organization and an egocentric metric that translates synaptic strengths to perceived changes in allocentric position [17].

The results also point out the importance of a recent criticism in the field that novel technical approaches in the study of neural circuits have neglected the use of behavioral assays [58]. In our study, behavior was not only a critical measure of the effects of 6 Hz optogenetic stimulation, it also interacted with our electrophysiological measures and entrainment efficacy. This interaction between stimulation and cognition has also been found in patients with epilepsy. The effect of electrical stimulation in memory areas of the brain on the acquisition of word information during a verbal memory task was determined by current brain state [59]. If specific structures were determined to be in a low encoding state, stimulation improved acquisition. If already in an encoding state, stimulation was detrimental to acquisition. Therefore, stimulation efficacy, to the benefit or detriment of cognition may depend on the task-mediated state of the circuit [60, 61].

Taken together, our results show that hippocampal inputs may compete for influence at local synapses in a manner that is mediated by local microcircuits and cognitive demand. This competition is not only found at the level of synaptic current, reflected in the properties of LFP theta oscillations, but also at the level of oscillatory patterns in CA1 pyramidal cell action potentials. Optogenetics not only has the potential to influence neural circuits, it can reveal changing dynamics in specific components of the circuit that underpin cognition. A key element for revealing these dynamics was the titration of cognitive demand in tasks that were behaviorally similar. This approach showed that cognitive demand significantly influences the entrainment efficacy of artificial theta inputs and that endogenous theta oscillations may be necessary for accurate self-localization. This reciprocal relationship between cognition, circuit state and entrainment efficacy is particularly important to consider as more studies begin to use optogenetic stimulation as a means of correcting circuit dysfunction in the hope of ameliorating cognitive deficits associated with temporal discoordination.

---

**Supplemental Figure 1:**
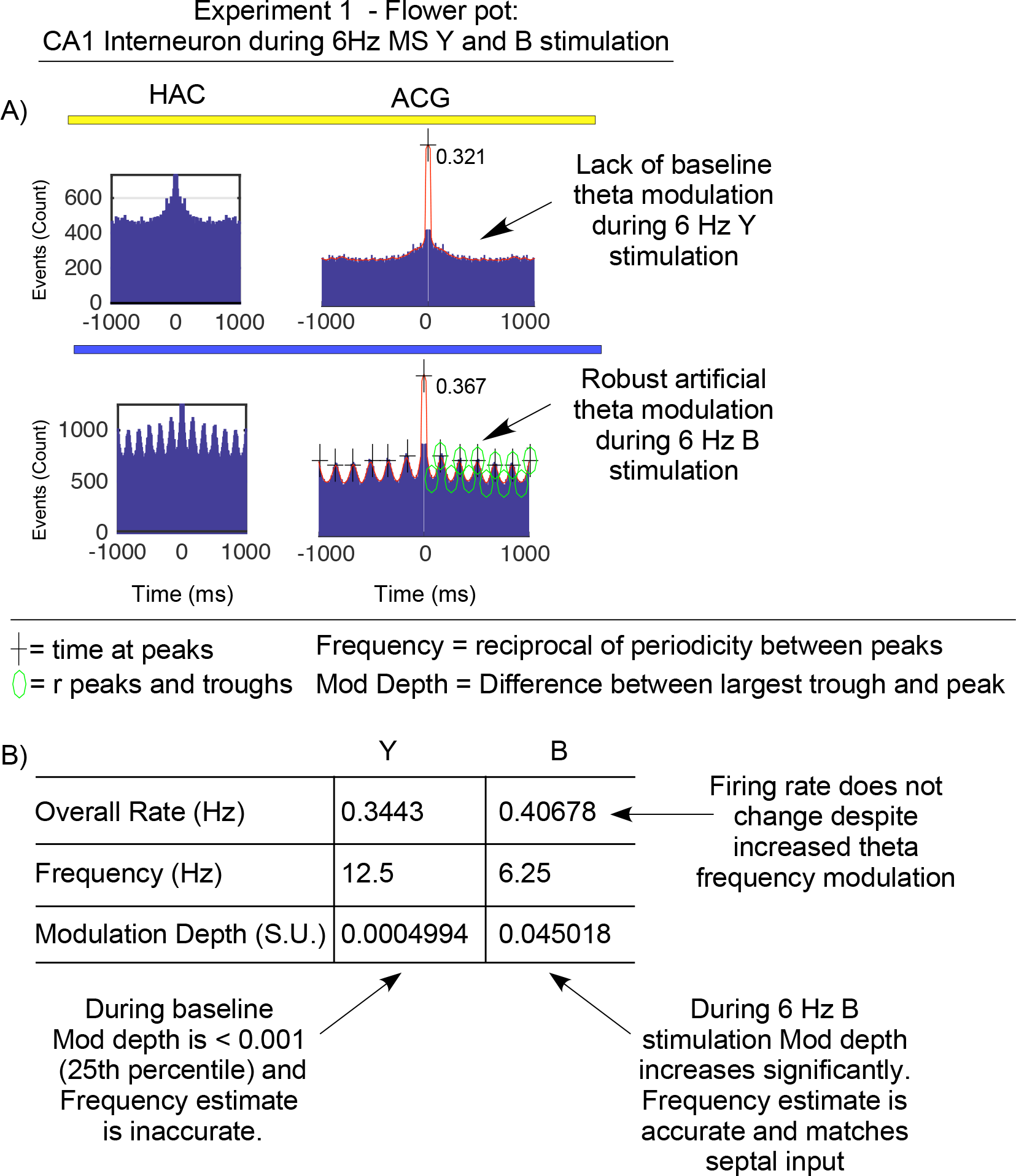
**A)** Example spike timing measurements during Experiment 1 illustrating modulation frequency and modulation depth results from a hippocampal interneuron during 6 Hz Y (Top) and B (Bottom) MS stimulation. For both conditions autocorrelation histograms (HAC at Left) and autocorrelograms (ACG at Right) are shown. Peak autocorrelation values are shown to the right of the peak ACG to provide scale for the Y-axis. The + symbols indicate the algorithm’s search for autocorrelation peaks to calculate periodicity. Green circles indicate the algorithm’s measurement of modulation depth between the largest trough and largest peak. Modulation frequencies tend to be high (i.e. > 9 Hz) when the modulation depth is low (i.e. < 0.001). **B)** Table illustrating the mean overall rate, modulation frequency and modulation depth of the example interneuron in A. While the cell’s firing rate remains constant, its modulation frequency and modulation depth are markedly altered to match the 6 Hz MS input signal during B stimulation.

**Supplemental Figure 2:**
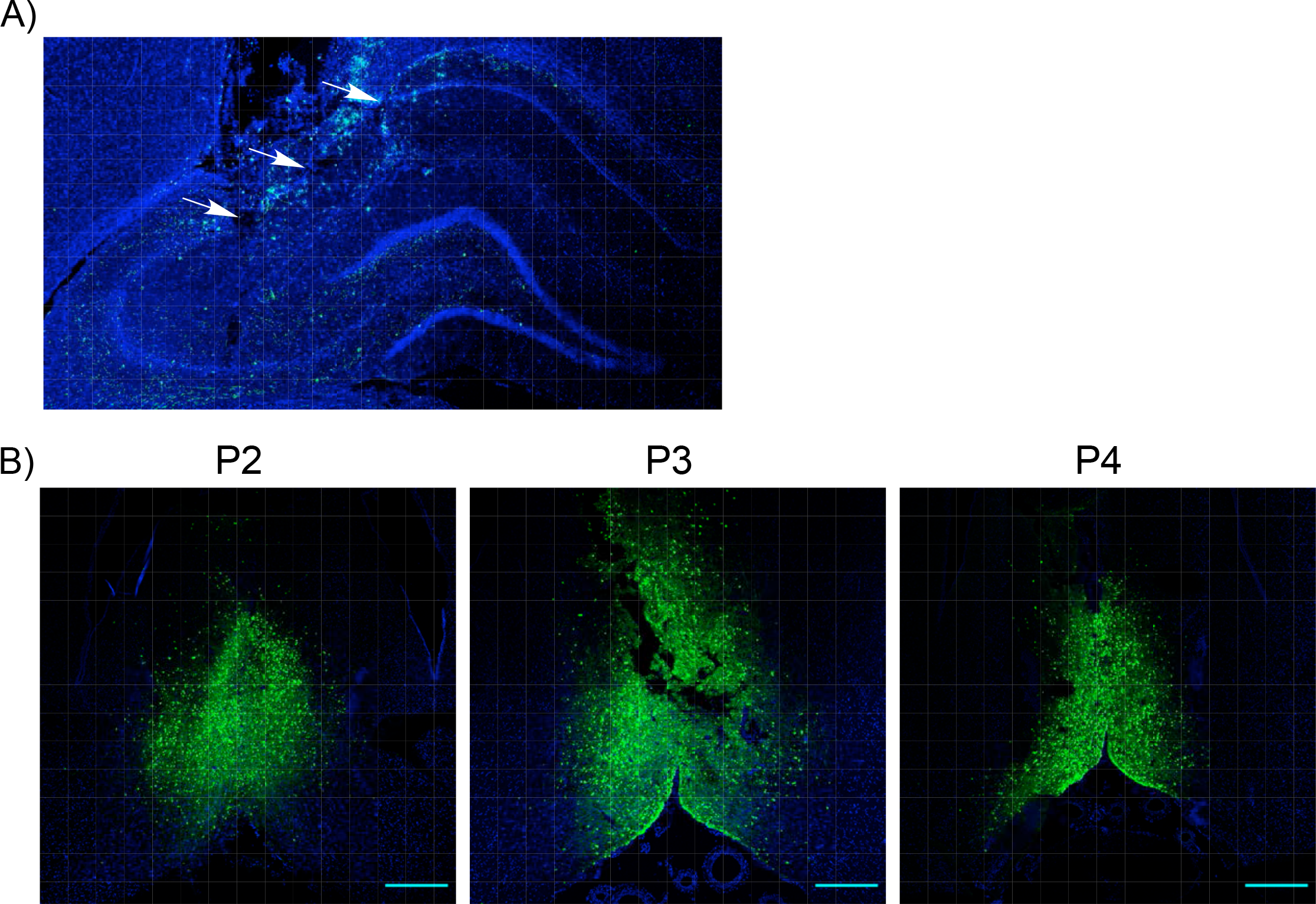
**A)** Recording tetrode tracks through the neocortex and CA1 of the dorsal hippocampus (White Arrows). Note heavy projection of transduced MS axons to the alveus layer and the dentate gyrus. **B)** Estimation of the volume of viral transduction in the MS for P2, P3 and P4. Estimated GFP expression volumes were consistent with the estimated total volume of the adult rat MS (mean = 1.69 ± 0.3 mm^3^). While the density of GFP expression was similar at the injection site in the MS across all rats, rat P2 exhibited less viral expression in the horizontal limb of the diagonal band of Broca (hDB) than Rats P3 and P4. Scale bar on lower right of each section = 500 μm. GFP expressions volumes for each slice in each rat are shown in Supp. Table 1. **C)** At Top Left - MS GAD stained GABAergic cell bodies (Red) in red co-localized with GFP (Green) and DAPI (Blue) (white arrows); MS Chat stained cholinergic cell bodies (Red) in red co-localized with GFP and DAPI (white arrows); MS VGLUT stain (Red) shows poor co-localization with GFP and DAPI at the soma. Bottom Left – 10x resolution image of a VGLUT (Red) stained hippocampal section with septal projection axons detected via GFP and hippocampal neurons detected via DAPI. Scale bar = 500 μm. Right – Three-dimensional rendering of 11 images taken at 2 μm steps of the section on the lower left (white box). Yellow puncta indicate the co-localization of VGLUT from MS axonal terminals and GFP in both str. pyramidale and str. oriens. Scale bar = 50 μm.

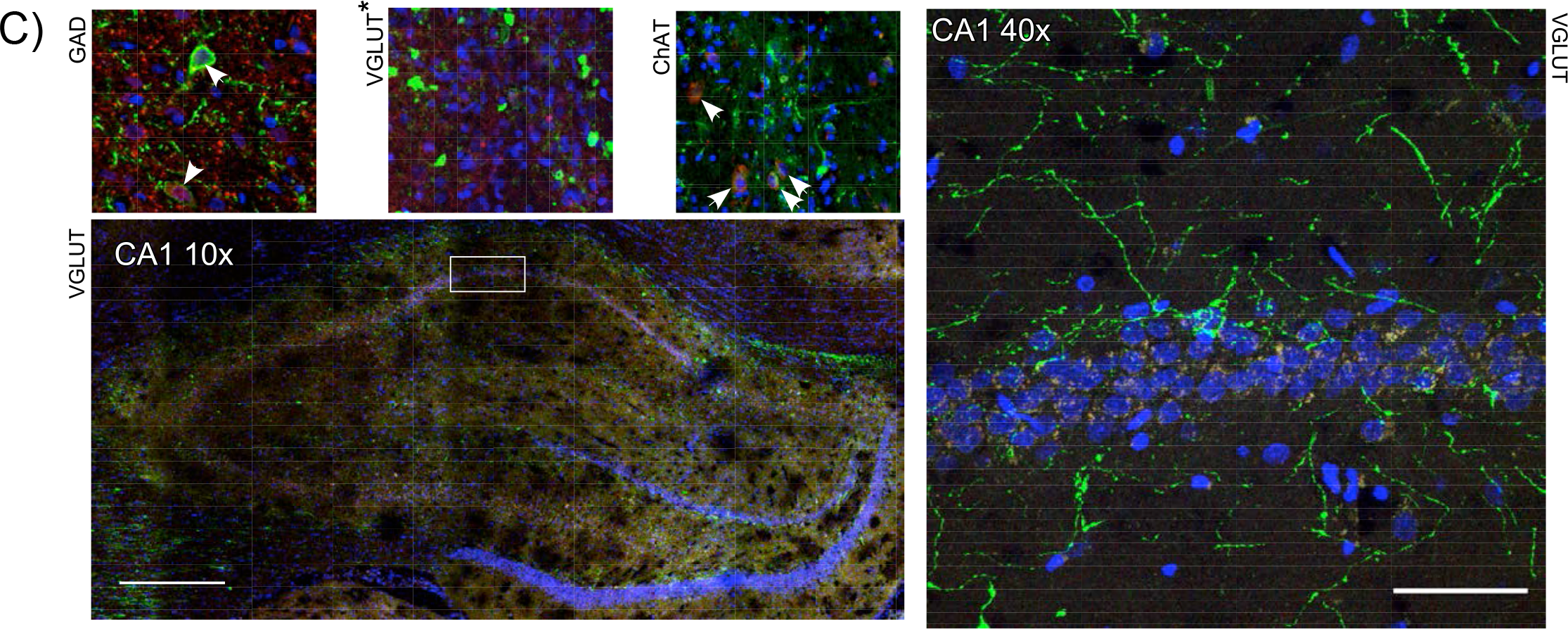

**Supplemental Figure 3:**
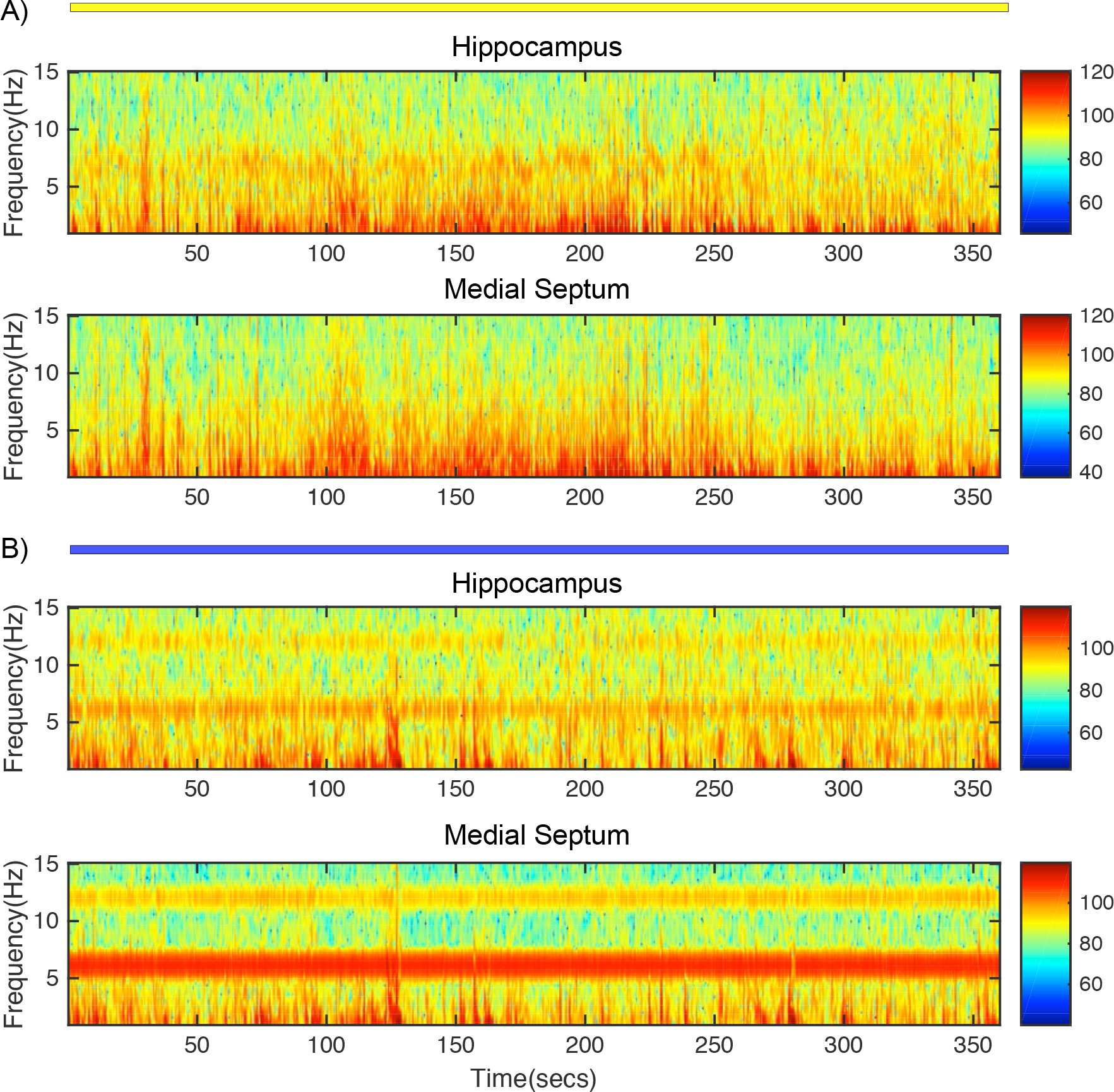
Experiment 1 - Full 6 min spectrogram data from the example shown in Figure 3. Results from both hippocampus (Top) and MS (Bottom) recordings during 6 Hz Y **(A)** and B **(B)** stimulation are shown. Color continuum at right indicates signal amplitude (A.U.). Endogenous theta signal in both regions is relatively weak as the rat is predominantly at rest. However, artificial theta generated during 6 Hz B MS stimulation is robust in this condition.

**Supplemental Figure 4:**
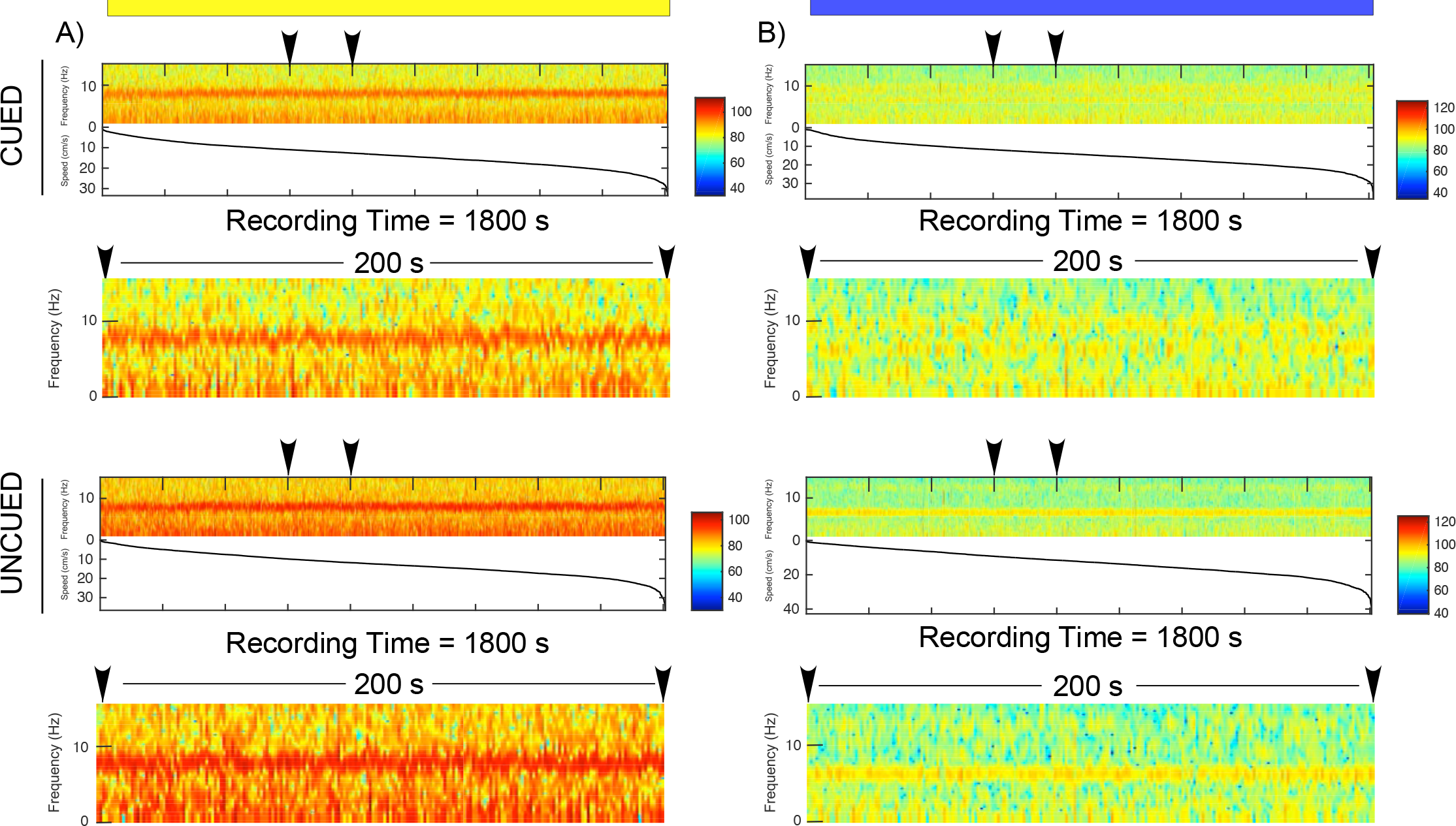
Experiment 2 - **A)** Speed-sorted spectrograms for CA1 theta oscillations during 6 Hz Y stimulation in the cued (Top) and uncued (Bottom) spatial accuracy tasks. Data corresponds to examples in Fig. 5A. Theta amplitude (Color bar at right in A.U.) and frequency increase linearly with speed. Black arrowheads indicate 200 s epochs of the speed-sorted spectrograms that have been expanded beneath each plot. **B)** As in A during 6 Hz B stimulation during the cued (Top) and uncued (Bottom) spatial accuracy tasks. During B stimulation in the cued task, endogenous theta frequency at approximately 9 Hz competes with the septal input frequency at 6 Hz. During B stimulation in the uncued version of the task, theta frequency predominately oscillates at 6 Hz.

**Supplemental Figure 5:**
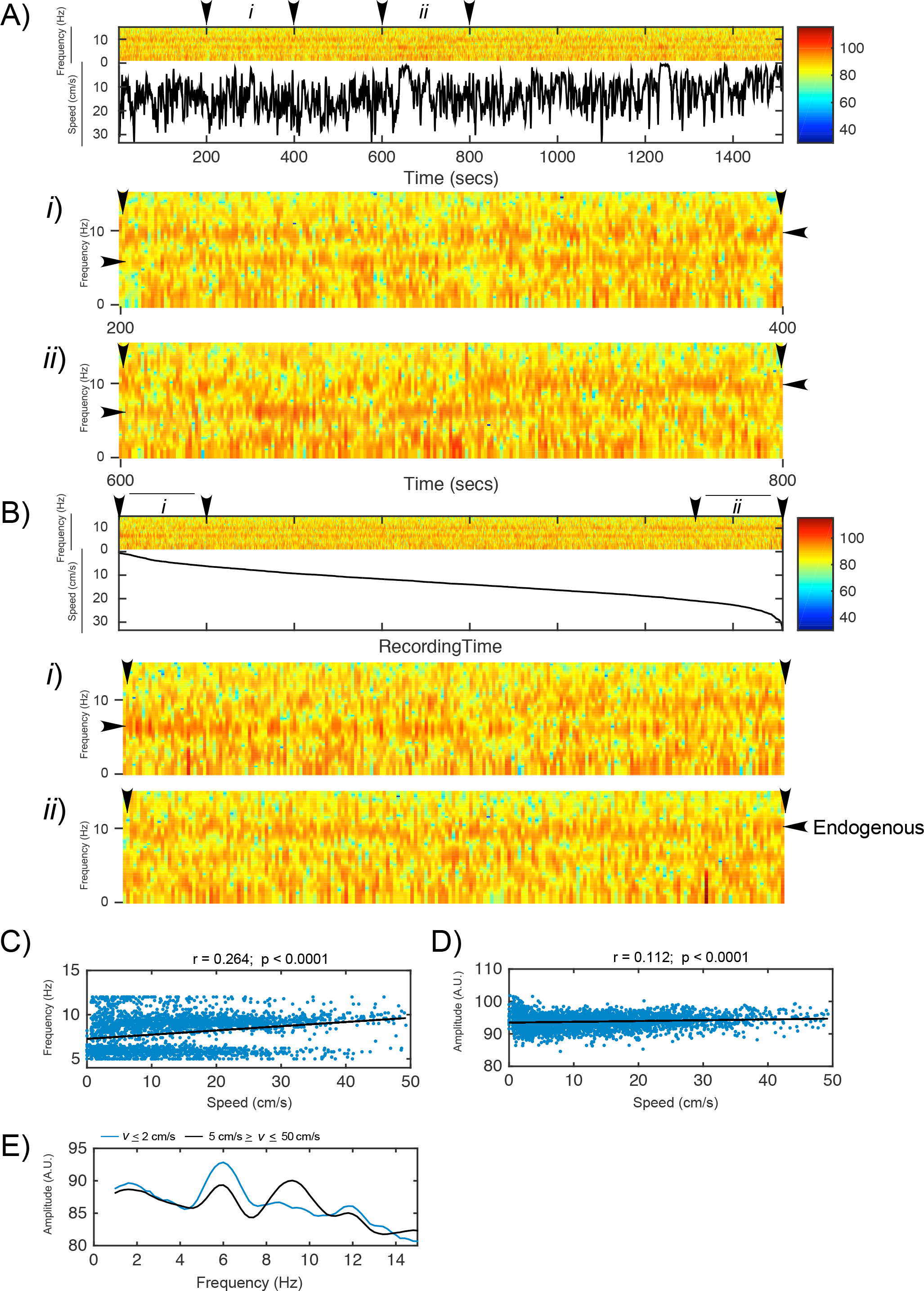
Experiment 2 - An additional example of frequency competition between endogenous and artificial theta for CA1 theta oscillations during 6 Hz B stimulation in the cued version of the spatial accuracy task. **A)** Spectrogram and animal speed for the full 30 min recording session (Top). Black arrowheads indicate approximately 200 s epochs that were selected for when the animal was primarily moving (*i*) or at rest (*ii*). These sections are expanded for visual inspection (Bottom). **B)** The same spectrogram data in A sorted by animal speed. Approximately 200 s of sorted data were selected at the extremes of animal speed when the animal was moving slowly (*i* = < cm/s) or quickly (*ii* = > 20 cm/s). These sections are expanded for visual inspection (Bottom). The data presented in A and B both indicate that hippocampal theta frequency more closely resembles artificial theta input during slow movements. During faster movements the amplitude of the endogenous theta frequency tends to be more dominant than the 6 Hz band. During intermediate speeds the theta signal tends to toggle between both frequencies. **C-D)** As in Fig. 5, the correlation between movement speed and the frequency and amplitude of each theta cycle (blue dots). The line of best fit (black line) and the corresponding correlation coefficients and p values are shown above each plot. **E)** Mean theta amplitude by frequency during slow (≤ 2cm/s = blue lines) and fast (≥ 5 cm/s = black lines) velocities (*v*) during Y and B MS stimulation is shown. The correlation between movement speed and the frequency and amplitude of each theta cycle (blue dots) for each condition is illustrated while the line of best fit (black line) and the corresponding correlation coefficients and p values are shown above each plot. Taken together these analyses indicate that in the cued session, while the rat is at rest or moving slowly (≤ 2cm/s), the frequency and amplitude at the septal input is more dominant. Beyond ≤ 2cm/s, the signal toggles between septal input and an endogenous theta signal. The amplitude of the endogenous theta signal tends to be larger and more dominant during faster animal speeds.

**Supplemental Figure 6:**
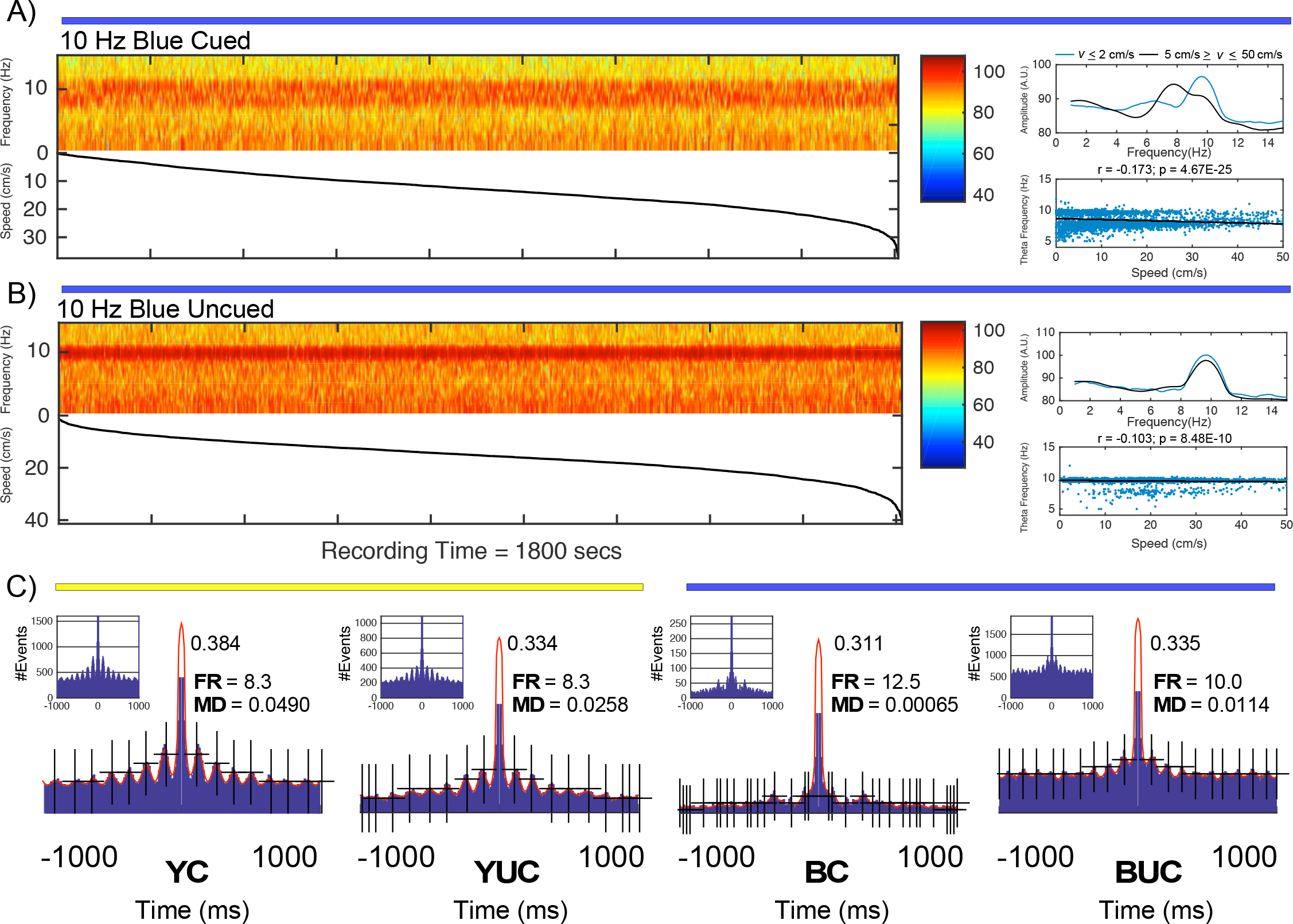
Experiment 2 - Sample LFP and spike timing results during B MS stimulation at 10 Hz during the cued and uncued spatial accuracy tasks. **A)** During the cued spatial accuracy task, speed sorted spectrogram (Left) as well as speed/theta frequency relationship (Right) as in Supp. Fig. 5. B stimulation causes an inversion of the speed/theta relationship. At slow movement speeds the majority of the theta band signal oscillates at 10 Hz. As the animal moves faster the theta signal toggles toward slower frequencies approaching mean theta frequency (~ 8 Hz). This results in a significantly negative speed/theta frequency relationship. **B)** Similar to the result found for 6 Hz stimulation, 10 Hz MS stimulation during the uncued version of the task results in theta band oscillations that more faithfully mirror the MS input frequency regardless of movement speed. **C)** Autocorrelation histograms (inset) and autocorrelograms illustrating theta modulation through inter-spike interval lag times for a CA1 pyramidal cell in each version of the spatial accuracy task during 10 Hz Y and B stimulation. Peak autocorrelation value, frequency (FR) and modulation depth (MD) are shown. During baseline sessions, the example cell is modulated at 8.3 Hz during the cued and uncued tasks. During B stimulation sessions, the cell is weakly modulated during the cued task but is robustly modulated at the septal input frequency during the task. Supp. Table 2: Mean + SE for speed and CA1 speed/theta properties by task condition across all 3 animals during 10 Hz B stimulation.

**Supplemental Figure 7:**
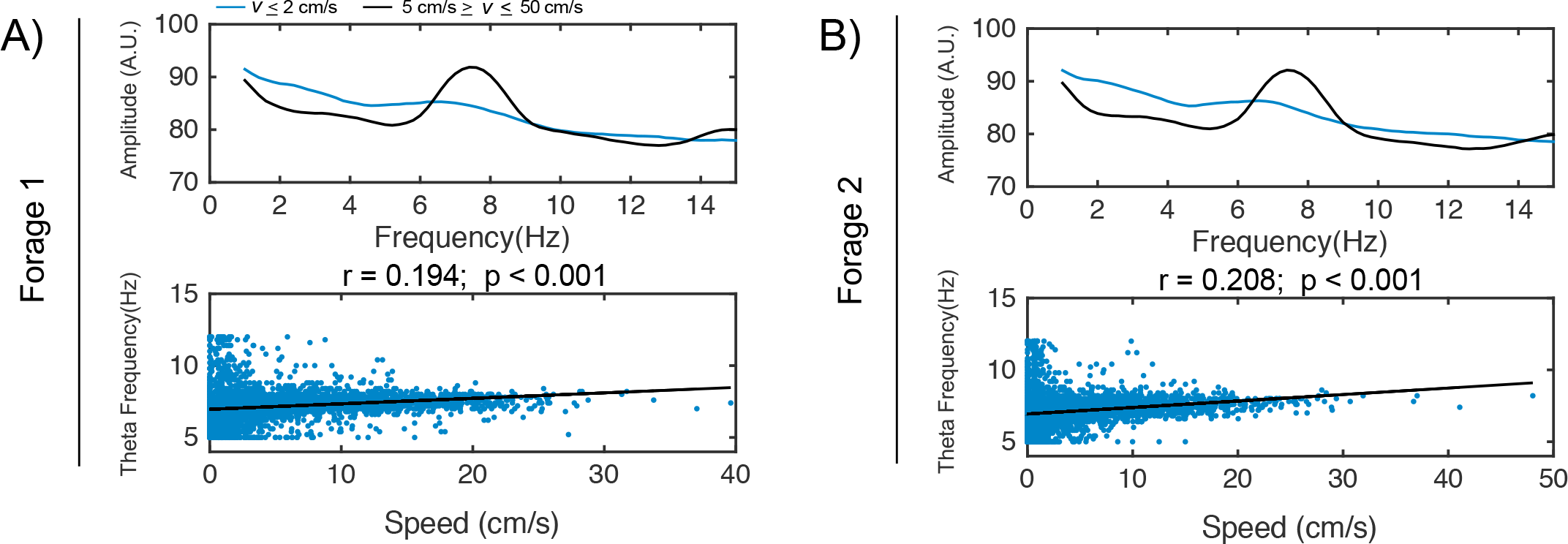
Experiment 2 - Examples of CA1 speed/theta properties from one animal during 2 consecutive food pellet foraging sessions (**A-B**) where theta frequency increases with movement speed in the absence of cognitive demand. Supplemental Table 3: Mean + SE for speed and CA1 speed/theta properties in each of the 2 foraging sessions across 3 animals. Peak theta frequency is faster during speeds ≥ 5 cm/s than speeds ≤ 2 cm/s. Mean speed/theta correlation in this condition is on par with the cued version of the spatial accuracy task during Y stimulation.

**Supplemental Figure 8:**
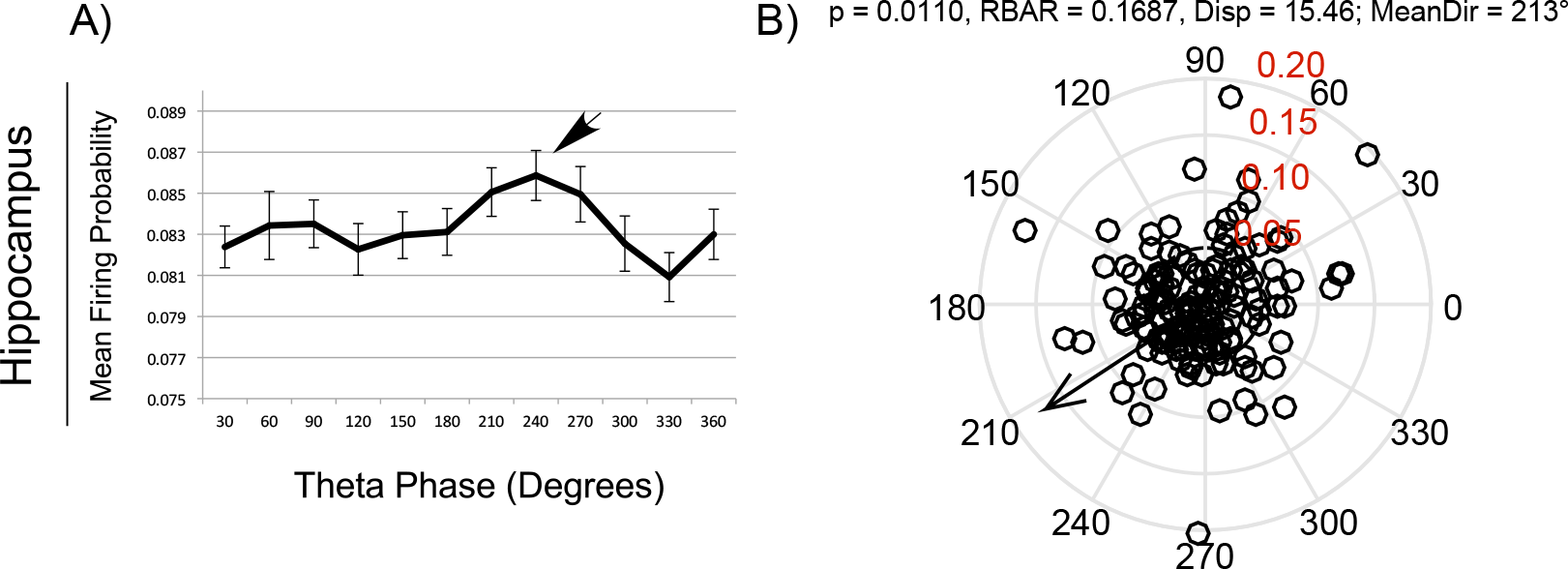
Experiment 2 - **A)** Firing probability mean + SE by theta phase for all CA1 pyramidal cells recorded from 3 animals (n =158) while foraging for food pellets. **B)** Pooled hippocampal theta phase preference circular statistics for the same cells represented in A. Each black circle represents the angle and magnitude of resultant theta phase vector (RBAR) for an individual pyramidal cell. The arrow indicates the angle and magnitude for the resultant population phase vector. RBAR levels for the circular plot are shown in red. The p-values, population RBAR value, dipersion level (Disp) and vector angle (MeanDir) are shown above the plot. The phase preference is diffuse across animals yet has a significant population vector and a similar level of dispersion as the pooled data for the YC condition in Fig. 5.

**Supplemental Figure 9:**
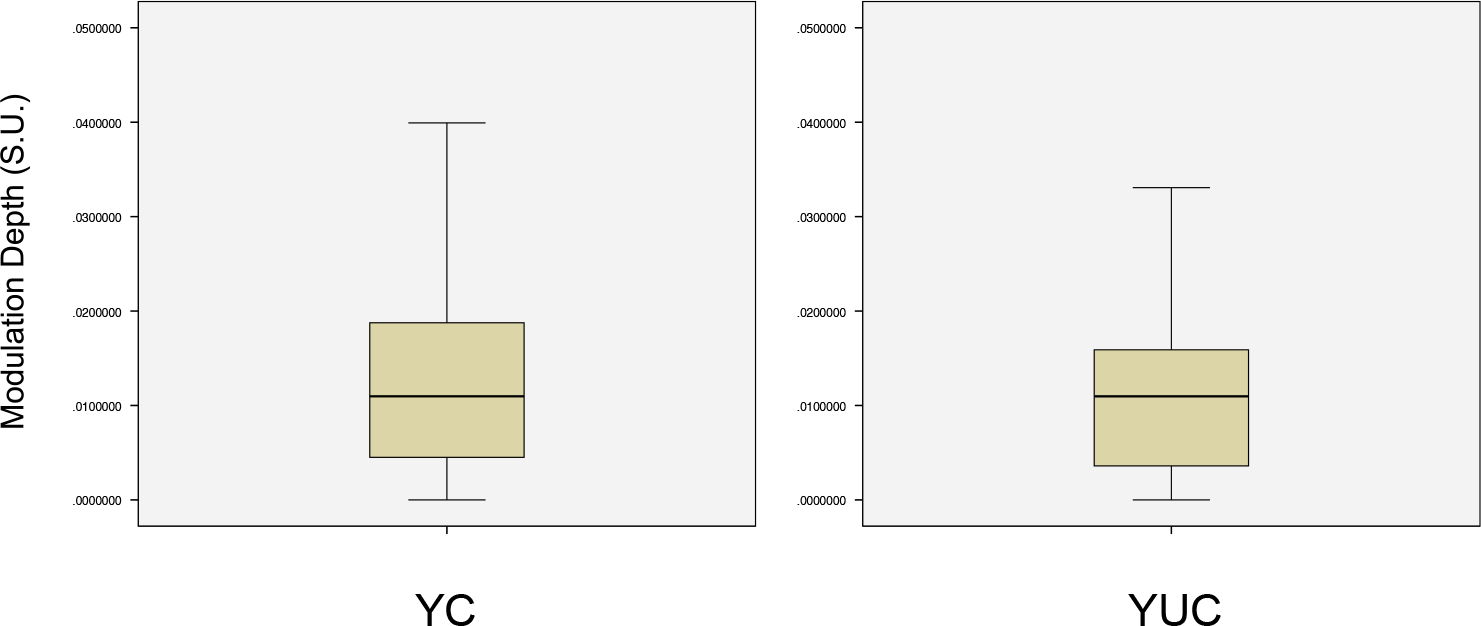
Experiment 2 - Boxplot figures for autocorrelation spike-timing analysis in Fig. 8. These boxplot data were used to establish Tukey’s hinge quartiles for the distribution of modulation depth for the cell population. The bottom quartile for the cells during YC (Left) was used to establish the lower cut-off for when cells were poorly modulated during BC and BUC sessions. The boxplot for the YUC session is also shown (Right) to demonstrate that the distribution was largely unchanged between YC and YUC.

**Supplemental Figure 10:**
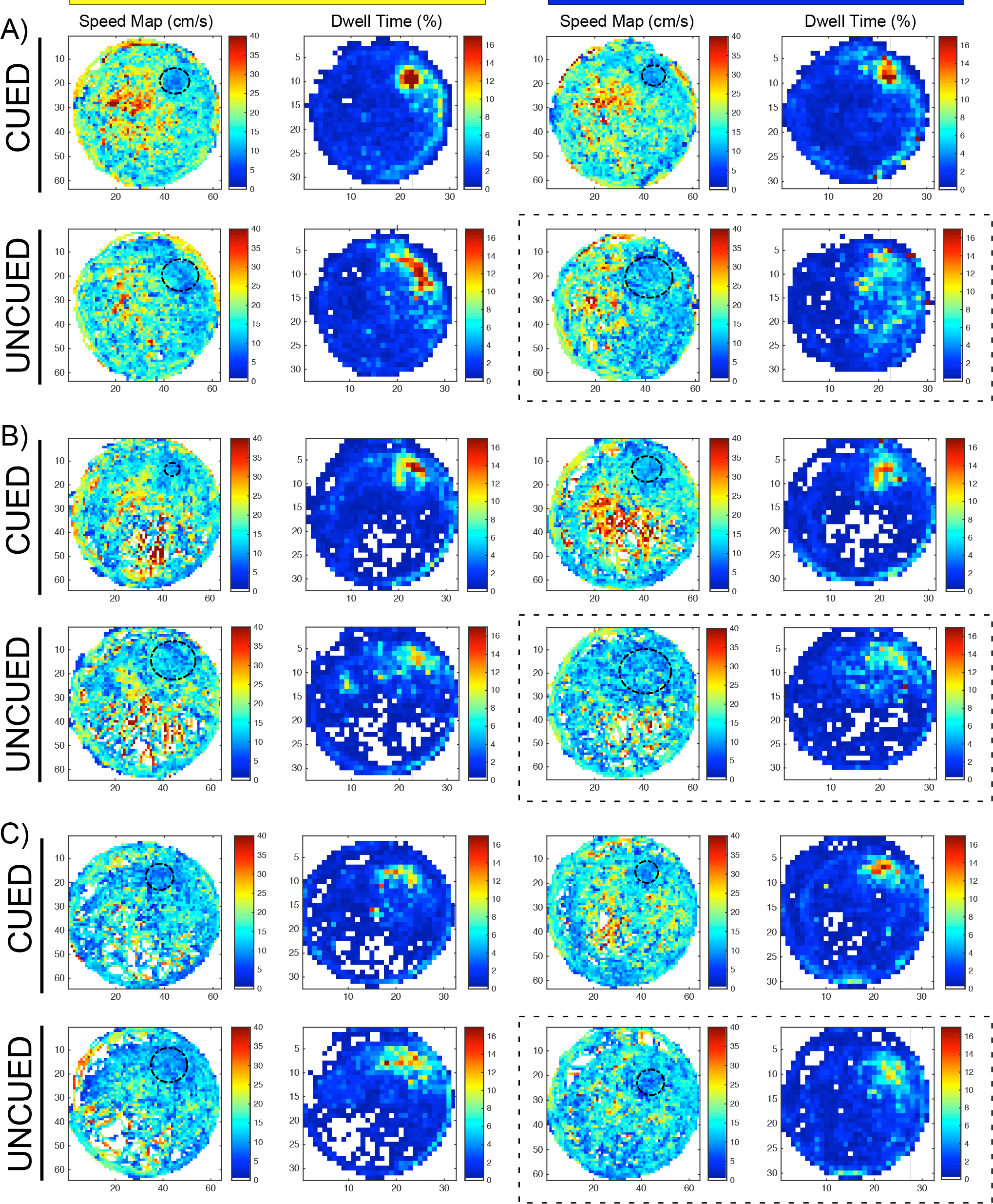
Experiment 2 - Three examples of spatial accuracy behavior are shown during each stimulation condition. **A-C)** Cued and uncued sessions are organized by row (Top and Bottom respectively) while 6 Hz Y and B sessions are organized by column (Left and Right Respectively). Each subsection consists of a speed map (cm/s) and dwell time map (% of overall time). These maps exhibit an inverse relationship as the rats cumulatively spend the majority of the session paused in the goal zone. Black dashed circles in the speed maps indicate the diameter of the rat’s mean search accuracy in the target quadrant. Relative to baseline performance during YC, 6 Hz B stimulation during the cued task had no effect on the rat’s ability to find the goal zone. In contrast, B stimulation during the uncued task (Dashed outline) generally resulted in the rat using a broader search pattern for locating the goal zone in the target quadrant (**A** and **B**) or consistently searching in the same off target location **(C)**.

**Supp. Table 1:**

| Animal ID | Area (mm<sup>2</sup>)<br>Anterior<br>Slice | Area (mm<sup>2</sup>)<br>Middle Slice | Area (mm<sup>2</sup>)<br>Posterior<br>Slice | Sum (mm<sup>2</sup>) | AP Distance<br>(mm) | Volume<br>(mm<sup>3</sup>) |
| --- | --- | --- | --- | --- | --- | --- |
| P2 | 0.175659 | 0.835616 | 0.665063 | 1.676338 | 1.2 | 2.0116056 |
| P3 | 0.671386 | 0.376186 | 0.286113 | 1.333685 | 1.2 | 1.600422 |
| P4 | 0.336196 | 0.807112 | 0.071398 | 1.214706 | 1.2 | 1.457647909 |

Mean = 1.69 mm<sup>3</sup> Standard Deviation = 0.30 mm<sup>3</sup>

**Supp. Table 2:**

|  | Speed (cm/s) | Speed/Theta r (S.U.) |  |
| --- | --- | --- | --- |
| BC | $12.88 \pm 1.72$ | $-0.035 \pm 0.057$ | |
| BUC | $13.92 \pm 1.85$ | $-0.042 \pm 0.034$ | |
|  | Peak Theta F (Fast) | Peak Theta F (Slow) | Mean Theta F |
| BC | $9.00 \pm 0.499$ Hz | $8.07 \pm 1.25$ Hz | $8.41 \pm 0.267$ Hz |
| BUC | $9.60 \pm 0.0$ Hz | $9.6 \pm 0.0$ Hz | $8.81 \pm 0.419$ Hz |
|  | Peak Theta Amp. (Fast) | Peak Theta Amp. (Slow) | Normalized Mean Theta Amp. |
| BC | $90.75 \pm 2.30$ | $92.28 \pm 3.65$ | $0.0600 \pm 0.00100$ |
| BUC | $91.95 \pm 2.98$ | $93.81 \pm 3.71$ | $0.0599 \pm 0.00087$ |

**Supp. Table 3:**

|  | Speed (cm/s) | Speed/Theta r (S.U.) |  |
| --- | --- | --- | --- |
| Forage 1 | 9.97 ± 2.87 | 0.225 ± 0.045 |  |
| Forage 2 | 9.18 ± 2.87 | 0.215 ± 0.043 |  |
|  | Peak Theta F (Fast) | Peak Theta F (Slow) | Mean Theta F (Fast) |
| Forage 1 | 7.60 ± 0.143 Hz | 6.60 ± 0.230 Hz | 7.56 ± 0.103 Hz |
| Forage 2 | 7.73 ± 0.150 Hz | 6.93 ± 0.241 Hz | 7.59 ± 0.103 Hz |
|  | Peak Theta Amp. (Fast) | Peak Theta Amp. (Slow) | Normalized Mean Theta Amp. (Fast) |
| Forage 1 | 95.55 ± 1.97 | 91.33 ± 3.09 | 0.0608 ± 0.000773 |
| Forage 2 | 95.48 ± 1.97 | 91.94 ± 3.11 | 0.0608 ± 0.000773 |

Author Contributions
J.M.B designed the experiments and analyzed the data; P.R.M and J.M.B collected data; M.L.K. analyzed histological and Immunohistochemistry data. All authors wrote the paper.

## Acknowledgements

We are very appreciative of the thoughts on analysis and the interpretation of our results provided by Dr. Kamran Diba, Dr. Colin Lever, Dr. Matt Wilson, Dr. Nathan Insel, Dr. Andre Fenton, Dr. Allan Gulledge and the assigned anonymous reviewers. We thank Andrew Alvarenga (GMW) for assistance with the custom design and manufacture of optical and electrophysiological implants. We thank Bruno Rivard, Rhys Niedecker for assistance with components of Figure 1. We are grateful for COBRE support from Sheryl White who greatly assisted with immunohistochemistry and from Todd Classon who assisted with imaging. We thank Pierre-Pascal Lenck-Santini, Sylvain Barriere and Sophie Sakkaki for assistance with Matlab code used for signal processing. We thank Mr. Daniel Mills for editing. We also acknowledge Neuralynx (Montana, USA) for dedicated technical support and Karl Deisseroth for supplying the adenovirus used in the study. This work would not have been possible without the advice of Dr. John Kubie during development and implementation of the spatial accuracy task. We declare no conflicts of interest. This work was supported by startup funds to JMB by UVM and by the NIH Grants NS108765 (GLH/JMB); NS108296 (GLH/JMB).

